# Developmental pleiotropy revealed by mosaic heterozygous *Snrpb* deletion underlies CCMS-like axial skeletal defects

**DOI:** 10.64898/2026.09.02.748635

**Authors:** Yanchen Dong, Eric Bareke, Marilou Paquin, Jennelle Smith, Sabrina Shameen Alam, Jacek Majewski, Loydie A. Jerome-Majewska

## Abstract

Cerebro-Costo-Mandibular Syndrome (CCMS) is a rare congenital disorder due to pathogenic variants in the core spliceosomal gene *SNRPB*. Affected individuals present with axial skeletal abnormalities, including cleft palate, micrognathia, posterior rib gaps, and a bell-shaped thorax. The molecular basis of these axial defects remains poorly understood, limiting the development of targeted interventions. To study the temporal and molecular requirements of SNRPB during axial development, we generated a mouse model of CCMS using an inducible Cre-lox system to conditionally delete *Snrpb*. Mosaic deletion of exons 2–3 of *Snrpb* after gastrulation resulted in the full spectrum of CCMS-like abnormalities, including micrognathia, posterior rib gaps and a reduced thoracic cavity. Despite normal somite morphology and patterning, transcriptomic analysis of E9.5 mutant somites revealed upregulation of p53 pathway genes and mis-expression of retinoic acid (RA) signaling components, consistent with reduced RA signaling. Alternatively spliced genes in *Snrpb* mutant somites were associated with post-transcriptional regulation, including chromatin modifiers. To test if RA pathway supplementation can rescue axial defects, we performed dietary RA supplementation; however, this failed to rescue mutant phenotypes, suggesting that p53 activation and chromatin dysregulation additionally contribute to disease pathogenesis. Together, these findings establish the first *in vivo* model of CCMS-associated axial skeletal abnormalities and indicate that *Snrpb* dysfunction disrupts the early specification of axial skeletal identity without overtly altering somite patterning.

## Introduction

Cerebro-Costo-Mandibular Syndrome (CCMS, OMIM#117650) is a rare congenital malformation associated with craniofacial and rib abnormalities. The defining features of CCMS are posterior rib gaps, and a bell-shaped thorax, which contribute to respiratory distress and feeding difficulties resulting in high mortality within the first year of life^1–5^. In a number of CCMS patients, pathogenic heterozygous variants that increase inclusion of an alternative exon 2 in the splicing factor *SNRPB* were identified^4,6^. Increased inclusion of this alternative exon, which contains a premature termination codon, is associated with reduced *SNRPB* levels in patient fibroblasts and is predicted to disrupt splicing of key genes essential for axial skeletal differentiation^4^.

*SNRPB* encodes for SmB/SmB’ that forms part of the heptameric ring necessary for biogenesis of the small nuclear ribonucleoprotein (snRNP) complexes of the major spliceosome, specifically U1, U2, U4 and U5^3,4^. As the major spliceosome is responsible for more than 99.5% of splicing reactions in humans^7^, how reduced levels of SNRPB give rise to tissue-specific effects in the axial skeleton remains an intriguing question. We recently found that SNRPB-mediated splicing regulates ER homeostasis and collagen secretion, consistent with a key role for splicing in cellular homeostasis^8^. Furthermore, work from our group and others indicates that SNRPB regulates splicing of factors important for survival, patterning, and differentiation of numerous cell types, including neural crest cells and osteoprogenitor-like cells^9,10^. In the latter, knockdown of *SNRPB* reduced survival and was associated with cell type-specific splicing changes^9^. Mouse embryos with constitutive heterozygous deletion of *Snrpb* exons 2–3 (including alternative exon AE2) produce 70% less *Snrpb* RNA and arrest at embryonic day (E) 9.5. Heterozygous deletion of these same exons of *Snrpb* in mouse neural crest cells (*Snrpb^ncc+/–^*) results in mis-splicing of genes associated with survival and craniofacial patterning^10^. In *Snrpb^ncc+/–^* mutant embryos, increased p53-mediated apoptosis and abnormal expression of growth factors such as *Fgf8* were associated with microcephaly and micrognathia^10^.

Here, we report that mouse embryos with mosaic heterozygous deletion of exons 2–3 of *Snrpb* shortly before E8.5 recapitulates CCMS-like skeletal abnormalities, including micrognathia, cleft palate, reduced thoracic cavity and rib gaps. Despite normal somite morphology and patterning, RNA sequencing of these embryonic precursors of ribs and vertebrae revealed enrichment of alternative splicing events in chromatin-modifying genes, along with altered expression of genes linked to p53 activity and retinoic acid (RA) signaling in mutants. We further show that while genes involved in retinoic acid metabolism are abnormally expressed, exogenous retinoic acid supplementation was not sufficient to rescue axial defects. Our findings indicate that reduced *Snrpb* drives developmental pleiotropy, disrupting cell survival, chromatin regulation, and RA signaling in morphologically normal somites. Our study indicates that axial skeletal identity is specified prior to overt somite segmentation and patterning and is vulnerable to levels of the core splicing factor SNRPB. Our data further suggest that somatic mosaicism for *SNRPB* should be investigated in clinically diagnosed CCMS patients without identified coding variants^5^.

## Results

### *Snrpb^tmx+/–^* mutants exhibit the full spectrum of CCMS-like abnormalities

To establish a model of CCMS, we first identified the embryonic stage at which tamoxifen-induced heterozygous deletion of *Snrpb* exons 2–3 produces CCMS-like abnormalities. At all stages, control littermates that also received tamoxifen (*Snrpb^tmx+/+^*), comprising of *Snrpb^+/+^; ER-CRE^tg/+^, Snrpb ^+/+^*; *ER-CRE ^+/+^,* and *Snrpb ^L/+^*; *ER-CRE ^+/+^* genotypes, were used for comparisons. Tamoxifen-induced deletion of *Snrpb* at E8.0 resulted in embryonic lethality by E12.5, while deletion induced after E8.5 failed to recapitulate CCMS-like axial skeletal abnormalities at E14.5, except for microcephaly (Supplementary Note; Table 1; Supplementary Fig. 1; Supplementary Data 1). In contrast, tamoxifen-induced *Snrpb* exons 2–3 deletion two hours before E8.5 (*Snrpb^tmx+/–^*) resulted in the full spectrum of CCMS-like axial defects (Table 2; Figure 1A). Thus, we conclude that wildtype levels of SNRPB are essential after gastrulation for normal axial development.

**Figure 1.**
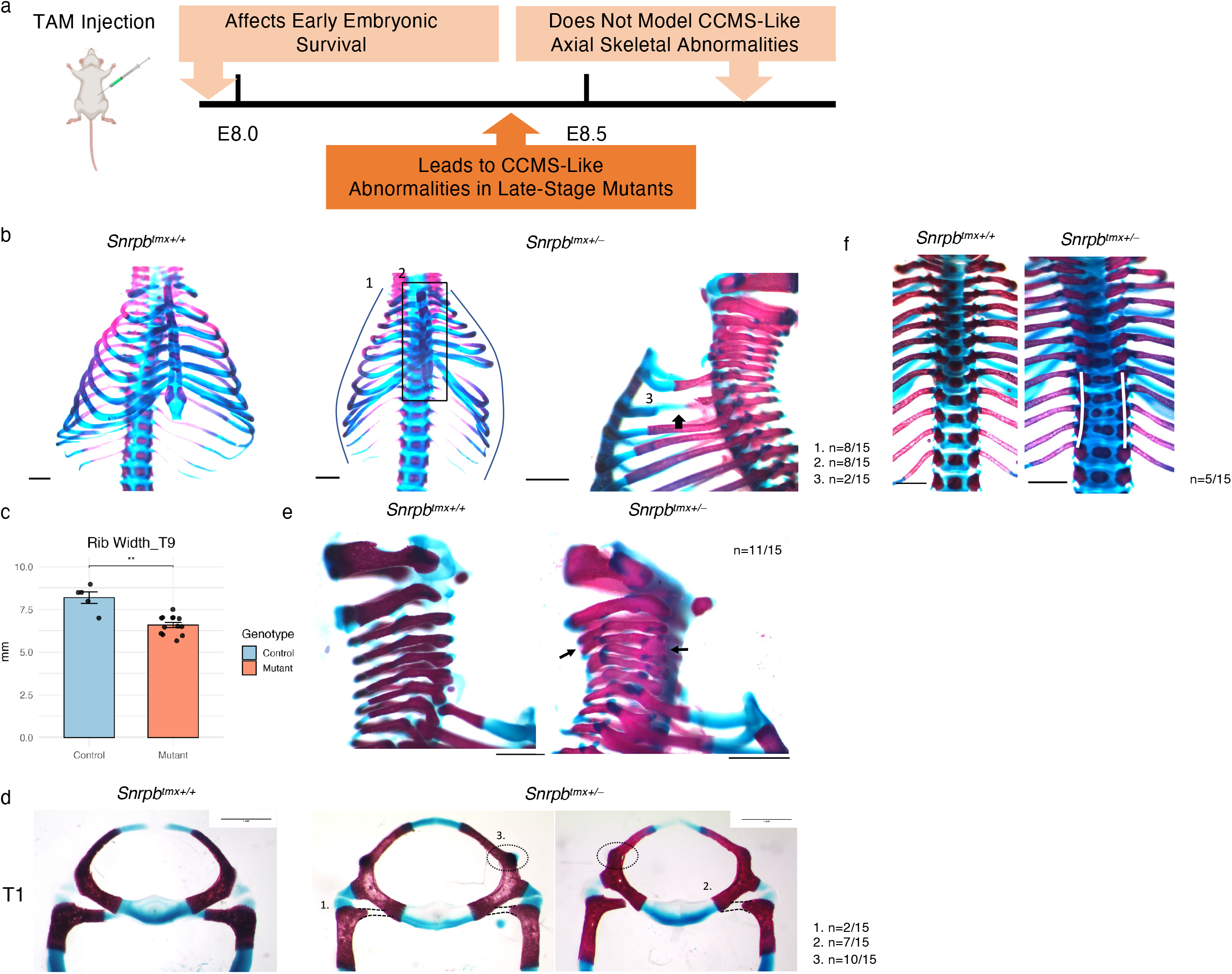
Rib and vertebral abnormalities in E17.5 *Snrpb^tmx+/–^* mutants. (a) Schematic illustrating the outcome of tamoxifen-induced *Snrpb* deletion at distinct embryonic stages. (b) Representative skeletal preparations showing rib abnormalities in mutants, including 1) a bell-shaped thorax (n=8/15), 2) asymmetric rib fusion to the sternum (n=8/15), and 3) posterior rib gaps (arrowheads; n=2/15). (c) Bar plot of T9 rib width in control and mutant embryos. Error bars represent standard error of the mean. Two-tailed Student’s t-test, **p<0.01. (d) Representative skeletal preparations showing anomalous rib– vertebra insertions, including 1) bilateral absence of the T1 rib head (n=2/15), 2) unilateral absence of the T1 rib head (n=7/15), and 3) ectopic tubercles on the T1 vertebra (n=10/15). (e) Representative skeletal preparations showing cervical vertebral fusions (arrowheads; n=11/15). (f) Representative skeletal preparations showing fusion of thoracic vertebrae resulting in scoliosis-like spinal curvature (n=5/15). Scale bars, 1mm.

**Table 1.**
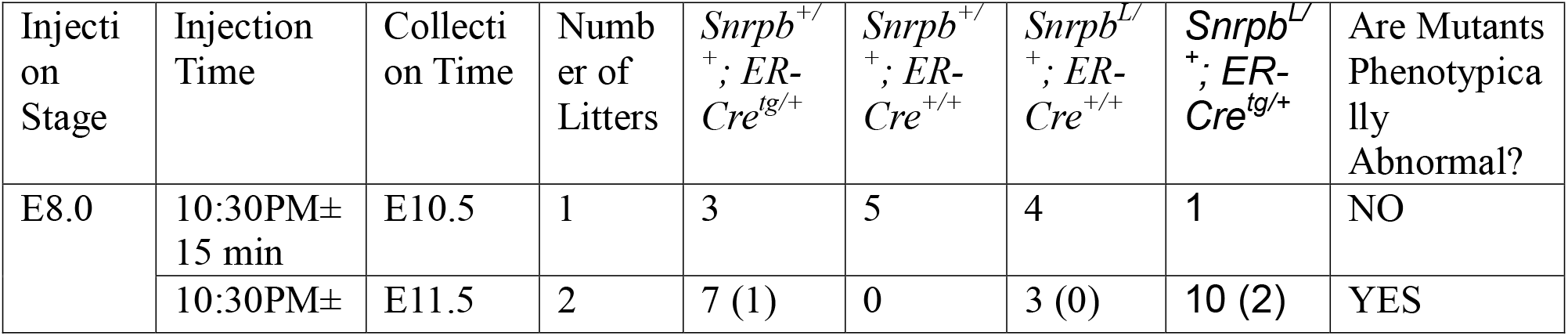
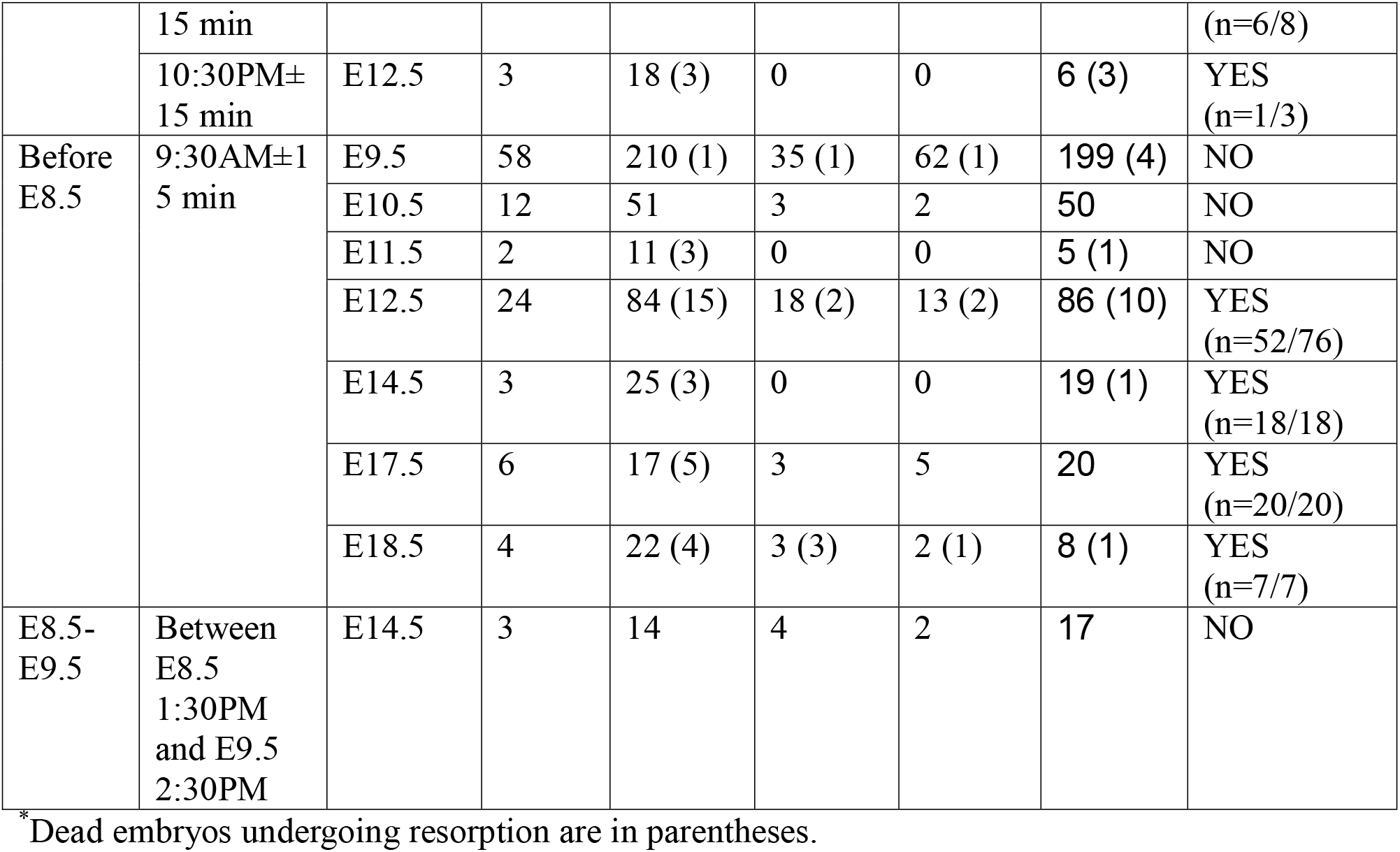
Summary of Tamoxifen Injection Times Tested, and Embryos Collected at Each Corresponding Time Point.

**Table 2.** CCMS Phenotypes Modeled in E17.5 *Snrpb^tmx+/–^* Mutants.

|  | CCMS (OMIM #117650) | Wnt1-Cre2 Model | TAM-Inducible Model |
| --- | --- | --- | --- |
| Head &<br>Neck | Microcephaly | ☐ | ☐ |
|  | Severe micrognathia | ☐ | ☐ |
|  | Malar hypoplasia |  |  |
|  | Cleft palate | ☐ | ☐ |
|  | Short palate |  |  |
|  | High-arched palate |  |  |
| Chest | Bell-shaped thorax |  | ☐ |
|  | Small thorax |  | ☐ |
|  | Rudimentary rib |  |  |
|  | Anomalous rib insertion to vertebrae |  | ☐ |
|  | Absent twelfth rib |  |  |
|  | Posterior rib gap defects |  | ☐ |
| Skeletal | Scoliosis |  | ☐ |
|  | Sacral fusion |  |  |

As appendicular defects in *Snrpb^tmx+/–^* embryos have been previously described^8^, we focus here on the axial phenotypes. At E14.5, cartilage preparations revealed microcephaly (two-tailed Student’s t-test, p < 0.0001) and a smaller, curved Meckel’s cartilage (two-tailed Student’s t-test, p < 0.0001; n=5/14) that was separated from the middle ear structures (n=10/14; Supplementary Fig. 2a-b). At E17.5, skeletal preparations confirmed microcephaly (two-tailed Student’s t-test, p < 0.01), micrognathia (two-tailed Student’s t-test, p < 0.0001), and abnormally attached middle ear ossicles at the proximal end of Meckel’s cartilage (n=14/15), and additionally revealed cleft palate (n=12/15; Supplementary Fig. 3a-d).

Importantly, *Snrpb^tmx+/–^* mutants also exhibited vertebral and rib malformations, hallmark features of CCMS. Cartilage preparations revealed rib defects, including asymmetric rib fusion to the sternum (n=7/14), false rib fusions (n=9/14), and reduced thoracic size (two-tailed Student’s t-test, p < 0.001; Supplementary Fig. 2c-d) in E14.5 mutant embryos. At E17.5, skeletal preparations confirmed reduced thoracic size, evidenced by a bell-shaped thorax (n=8/15; Figure 1b) and significantly reduced rib width (two-tailed Student’s t-test, p < 0.01; Figure 1c). Additional rib anomalies included posterior rib gaps (n=2/15; Figure 1b), abnormal rib attachment to the sternum (n=9/15; Figure 1b), missing rib heads (n=9/15; Figure 1d), and ectopic tubercles on the T1 vertebra (n=10/15; Figure 1d). Vertebrae abnormalities were also observed, including cervical vertebrae fusion (n=11/15; Figure 1e), thoracic scoliosis (n=5/15; Figure 1f), and elongated centrum with an abnormally prominent neural spine (n=14/15; Supplementary Fig. 4). Together, these findings demonstrate that *Snrpb* deletion shortly before E8.5 recapitulates key features of CCMS.

### Mosaic *Snrpb* loss uncouples axial skeletal defects from normal somite formation and sclerotome patterning

To determine how soon after tamoxifen injection Cre activity, thus *Snrpb* deletion, was induced, we introduced the mT/mG reporter into our mutant line. Embryos collected 10 hours after tamoxifen administration (3 litters; somite range: 4–14) showed mosaic GFP expression, indicating Cre activity, along the rostrocaudal axis in both control and mutant embryos (Figure 2a). In embryos of both genotypes, mosaic GFP expression was consistently found along the rostrocaudal axis, including in somites, presomitic mesoderm, and tailbud at 14.5 (3 litters; somite range: 7–17) and 24 hours post-injection (3 litters; somite range: 13-22; Figure 2a). This mosaic expression was maintained at 36 and 48 hours post-injection (Supplementary Fig. 5). These data indicate that *Snrpb^tmx+/–^* mutant embryos undergo mosaic deletion along the entire rostrocaudal axis, including in the craniofacial region, presomitic mesoderm, somites, and tailbud.

**Figure 2.**
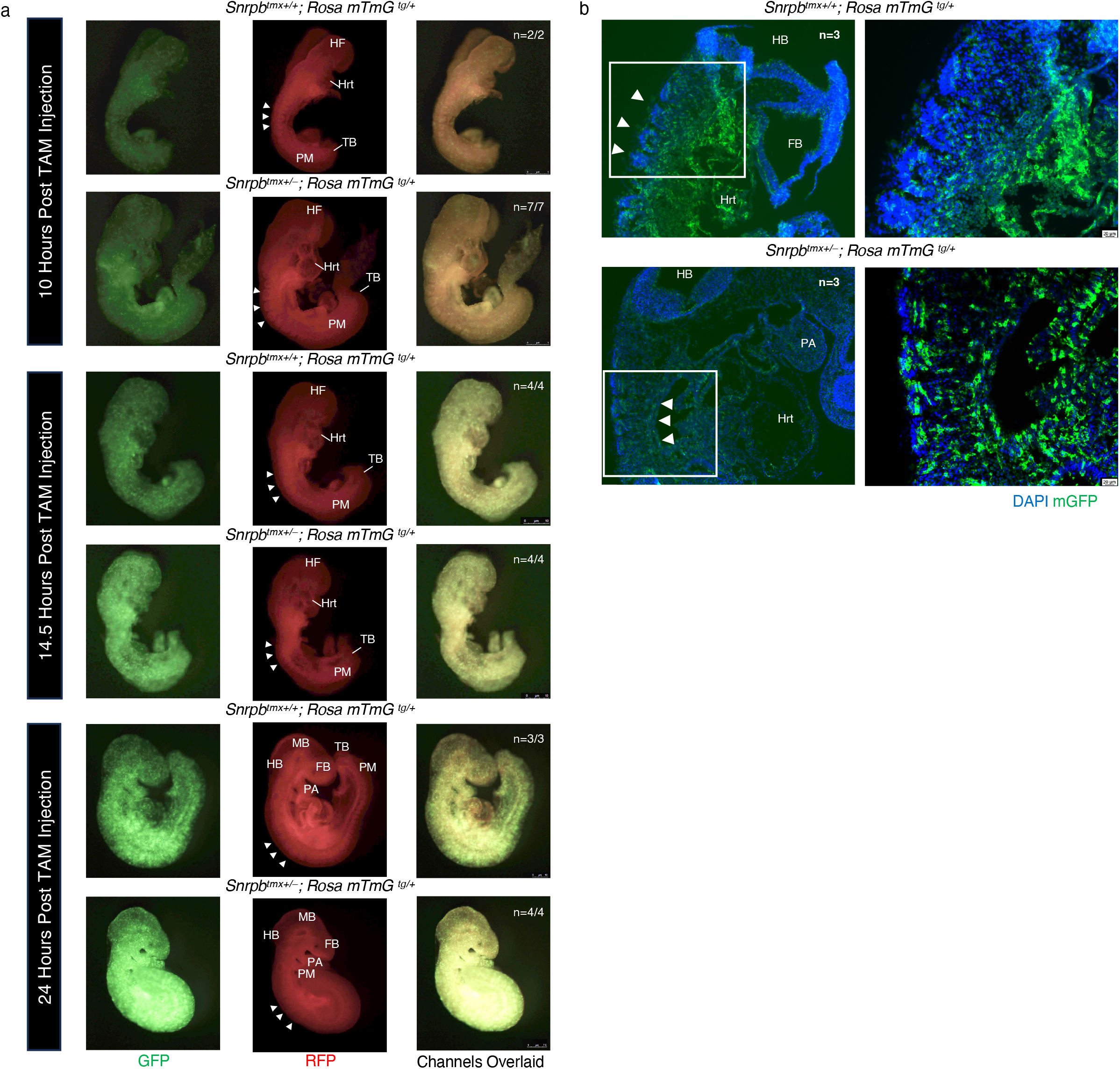
Mosaic *Snrpb* deletion along the rostrocaudal axis in *Snrpb^tmx+/–^* mutants. (a) Representative fluorescence images of control and *Snrpb^tmx+/–^*mutant embryos carrying the *Rosa*26-mTmG reporter at 10, 14.5, and 24-hours post-tamoxifen injection at E8.5, showing mosaic GFP expression along the rostrocaudal axis. (b) Representative sagittal sections of control and *Snrpb^tmx+/–^*mutant embryos carrying the *Rosa*26-mTmG reporter at E9.5, showing GFP-positive cells in somites. HF, headfold; Hrt, heart; PM, presomitic mesoderm; TB, tailbud; FB, forebrain; MB, midbrain; HB, hindbrain; PA, pharyngeal arch. Arrowheads indicate somites.

The sclerotome that differentiates to form the vertebral bodies, intervertebral discs, and ribs is derived from the ventral region of somites, which forms bilaterally in a rostro-caudal sequence between E8.0 and E11.5^11–16^. To determine if somites form in *Snrpb^tmx+/–^*mutant embryos, we collected embryos from E9.5 to E12.5. Prior to E12.5, *Snrpb^tmx+/–^*mutant embryos (n=249) were indistinguishable from controls (n=368) with no apparent differences in somite number or size. Furthermore, although 68% of E12.5 *Snrpb^tmx+/–^* mutant embryos (n = 52/77) exhibited a triangular forebrain, reduced frontonasal prominence, and shortened limbs (Supplementary Fig. 6a) morphological defects were not found in the trunk. Using *Snrpb* mutant embryos carrying the mT/mG reporter we confirmed that GFP-positive, *Snrpb* mutant, cells were present specifically within the somites. Sagittal sections of E9.5 control (n=3/3) and mutant (n=3/3) embryos exhibited a mosaic distribution of GFP-positive cells along the dorsoventral aspect of somites, including in the ventral half of each somite, which undergoes epithelial-to-mesenchymal transition (EMT) to form sclerotome^17^ (Figure 2b). To determine whether sclerotome specification and rostrocaudal somite polarity were established normally, we examined the expression of *Pax1, Uncx4.1,* and *Tbx18* in E9.5 embryos. The expression patterns of *Pax1*, a sclerotome marker, and the rostrocaudal polarity markers *Uncx4.1* and *Tbx18* were comparable to those observed in control embryos (Supplementary Fig. 6b-c), indicating that these processes were not overtly affected. Thus, we conclude that *Snrpb* mutant cells contribute to somites that are properly patterned to form the sclerotome.

### P53 activity is transiently increased in *Snrpb* anterior somites and is associated with cell-death during somite re-segmentation

Since *Snrpb* is required prior to E8.5 and *Snrpb^tmx+/–^* embryos displayed normal somite formation and patterning between E9.5 and E11.5, we concluded that *Snrpb* is required for expression or splicing of transcripts important after sclerotome specification and/or resegmentaton. To identify the differentially spliced and/or expressed transcripts in the tailbud, paraxial mesoderm, and/or somites (n=3 per genotype) we used bulk RNAseq. Using an adjusted p-value < 0.05 and a fold change > 1, we identified 114 differentially expressed genes (DEGs) in *Snrpb^tmx+/–^*mutant samples. Intriguingly, 4.3-fold more genes were upregulated, with 93 upregulated and 21 downregulated in mutant samples (Figure 3A; Supplementary Data 2). Gene ontology analysis revealed no significantly enriched pathways among downregulated genes (Supplementary Fig. 7a), whereas upregulated DEGs showed enrichment for signal transduction by p53 class mediators, suggesting activation of cellular stress pathways in *Snrpb* mutant somites (Figure 3b).

**Figure 3.**
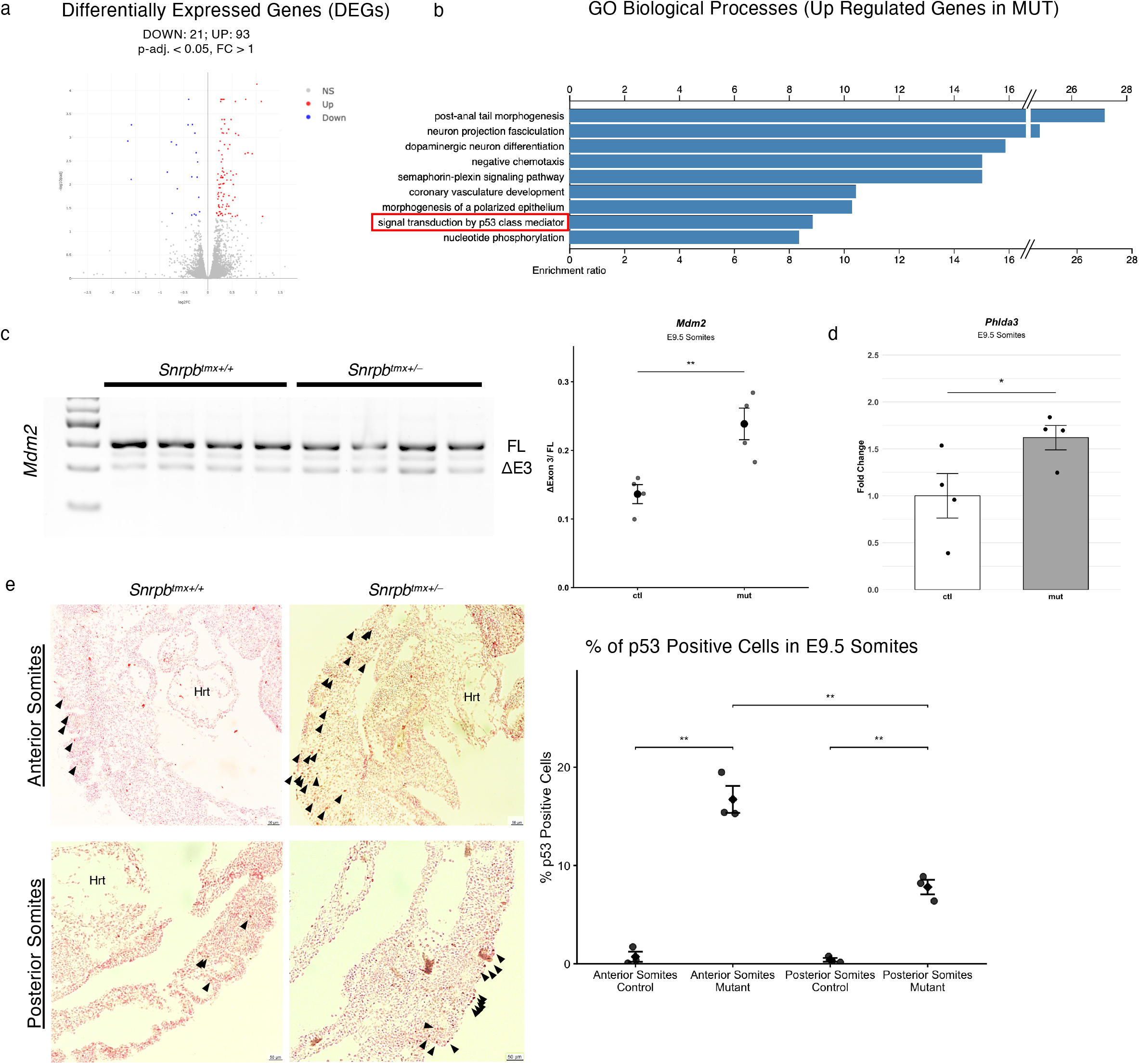
p53 pathway activation in E9.5 *Snrpb^tmx+/–^* mutant somites. (a) Volcano plot showing differentially expressed genes in *Snrpb^tmx+/–^* mutant samples relative to control from bulk RNA-seq performed on three pooled control and three pooled mutant samples (four embryos per pool; 25–28 somite stage), with 93 upregulated and 21 downregulated genes (adjusted p < 0.05, fold change > 1). (b) Gene ontology biological process terms associated with differentially expressed genes in *Snrpb^tmx+/–^* mutants (FDR<0.05). (c) Representative RT–PCR gel showing alternative splicing of *Mdm2* exon 3 in E9.5 control and mutant samples, and dot plot (mean ± SEM) of *Mdm2* exon 3 skipping levels (one-tailed Student’s t-test, **p < 0.01). (d) Bar graph (mean ± SEM) showing *Phlda3* expression levels in E9.5 control and mutant samples (one-tailed Student’s t-test, *p < 0.05). (e) Representative immunohistochemistry sections showing p53-positive cells (arrowheads) in anterior and posterior somites of control and *Snrpb^tmx+/–^* mutant embryos at E9.5, and dot plot (mean ± SEM) showing the proportion of p53-positive cells between genotypes (one-tailed Student’s t-test, **p < 0.01). Hrt, heart.

We previously showed that increased skipping of exon 3 of *Mdm2* and exon 7 of *Mdm4* results in increased p53-associated cell death and contributed to craniofacial defects in embryos with *Snrpb* mutant neural crest cells^10^. Therefore, we first assessed whether p53 activity and cell death were increased during somitic re-segmentation and compartmentalization. In somites, paraxial mesoderm, and tailbud isolated from E9.5 embryos, we observed a significant increase in *Mdm2* exon 3 skipping compared to controls (one-tailed Student’s t-test, p < 0.01; Figure 3c), which was further supported by our RNA-seq data (FDR < 0.05, inclusion level difference = 0.094), whereas *Mdm4* exon 7 splicing was comparable between genotypes (Supplementary Fig. 7b). Additionally, among three p53 targets previously found to be increased in *Snrpb* mutant neural crest cells^10^, RNA-seq analysis identified *Phlda3* and *Ccng1* as significantly upregulated in mutant samples (adjusted *p* < 0.05, fold change > 1). By RT-qPCR, only *Phlda3* was found to be significantly increased (one-tailed Student’s t-test, *p* < 0.05; Figure 3d), whereas *Ccng1 and Trp53inp1 were* not changed (Supplementary Fig. 7c). We also examined embryos at E12.5, when morphological abnormalities were first observed, to determine whether these changes in *Mdm2*/*Mdm4* splicing and p53 target gene expression persist. At E12.5, RT-PCR of trunk tissues showed no significant difference in *Mdm2* or *Mdm4* splicing, nor any changes in expression of the three p53 target genes assessed (Supplementary Fig. 7d-e). Together, these findings indicate that p53 activation in *Snrpb^tmx+/–^* mutant embryos is transient and confined to somitic progenitors rather than their differentiating descendants, pointing to a stage-specific requirement for *Snrpb* in regulating p53 activity during axial skeletal development.

Using immunohistochemistry, we examined the distribution of nuclear p53 in anterior and posterior somites, defined relative to the first forelimb bud, of E9.5 embryos. A significant increase in cells with nuclear p53 was observed in both anterior and posterior somites of *Snrpb^tmx+/–^* embryos compared to controls (one-tailed Student’s *t*-test, *p* < 0.01; Figure 3e), consistent with an increase in p53 activity in mutants. As axial defects in *Snrpb^tmx+/–^* embryos predominantly affect structures derived from somites anterior to the first forelimb bud, including cervical and thoracic vertebrae and ribs^11,18^, we assessed whether p53 activation was similarly enriched in this region. We found a significantly greater proportion of cells with nuclear p53 in anterior somites (one-tailed Student’s *t*-test, *p* < 0.01), indicating that activation of the p53 pathway is most pronounced in region most affected by reduced *Snrpb* levels.

To determine whether this p53 activation was associated with cell death, as previously observed in *Snrpb* mutant neural crest cells^10^, we examined distribution of TUNEL positive cells in anterior somites of E9.5 embryos. The proportion of TUNEL-positive cells was comparable between control and mutant anterior somites (Figure 4a). Similarly, no differences in cellular proliferation were observed between genotypes in either anterior or posterior somites (Figure 4b), indicating that elevated p53 activity in E9.5 mutant anterior somites is not accompanied by increased cell death or altered proliferation. However, in morphologically abnormal E12.5 mutant embryos, the proportion of TUNEL-positive cells in anterior trunk tissues was significantly increased compared to controls (one-tailed Student’s t-test, p < 0.05; Figure 4c). Together with the elevated p53 activity observed at E9.5, these findings suggest that transient p53 pathway activation in anterior somite progenitors may contribute to cell death in their differentiating derivatives at E12.5.

**Figure 4.**
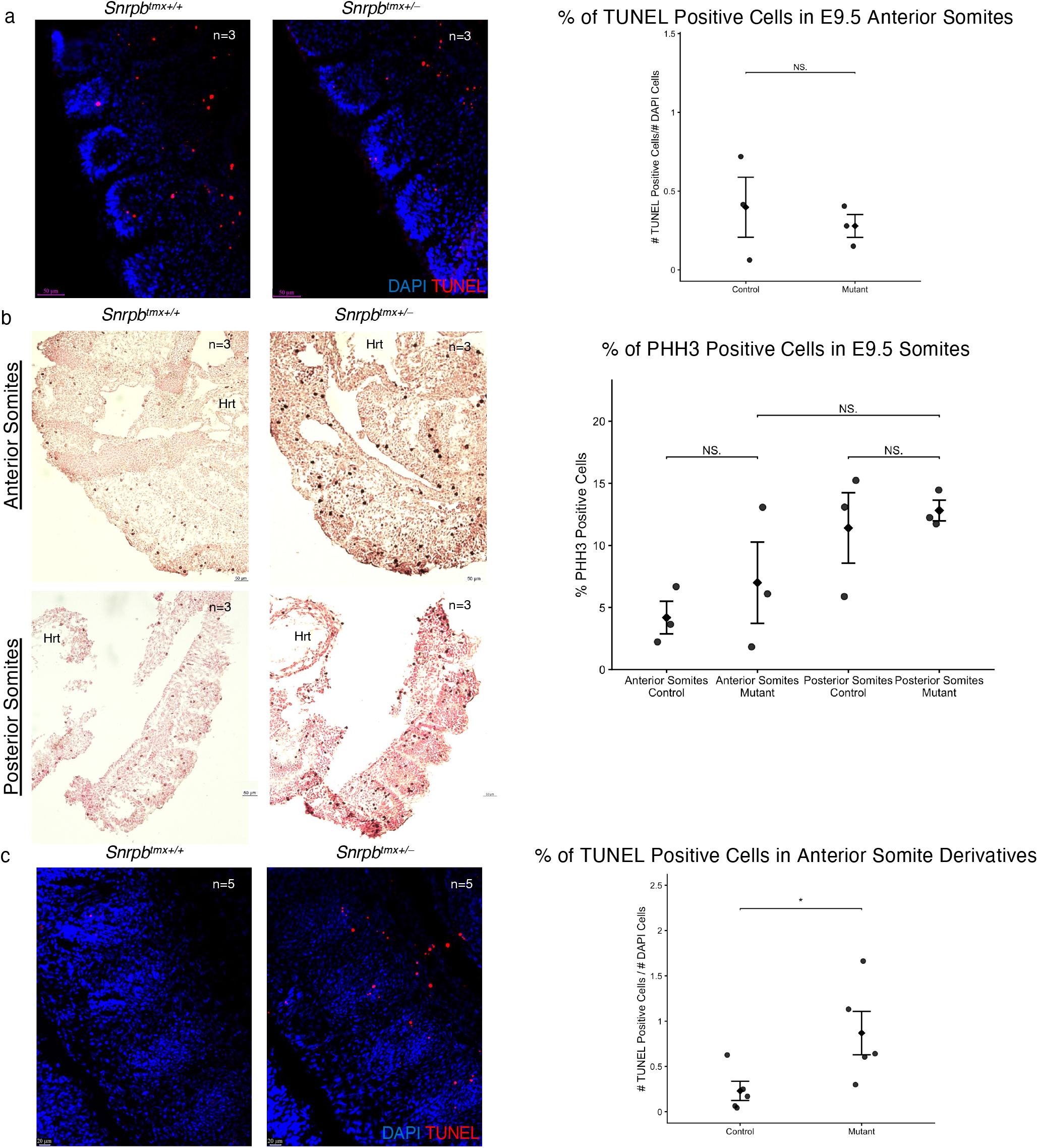
p53 pathway activation in *Snrpb^tmx+/–^* mutant anterior somites is not associated with immediate cell death or altered proliferation. (a) Representative TUNEL and DAPI-stained paraffin sections of anterior somites from E9.5 control and *Snrpb^tmx+/–^* mutant embryos, and dot plot (mean ± SEM) showing the proportion of TUNEL-positive cells between genotypes (one-tailed Student’s t-test, p > 0.05). (b) Representative phospho-histone H3 (PHH3) immunohistochemistry sections showing PHH3-positive cells in anterior and posterior somites of E9.5 control and *Snrpb^tmx+/–^*mutant embryos, and dot plot (mean ± SEM) showing the proportion of PHH3-positive cells between genotypes (one-tailed Student’s t-test, p > 0.05). (c) Representative TUNEL and DAPI-stained paraffin sections of anterior somite-derived tissues from E12.5 control and *Snrpb^tmx+/–^* mutant embryos, and dot plot (mean ± SEM) showing the proportion of TUNEL-positive cells between genotypes (one-tailed Student’s t-test, *p < 0.05). Hrt, heart.

### Retinoic acid signaling is perturbed in *Snrpb^tmx+/–^* mutant embryos

To identify mis-expressed transcripts or pathways important for vertebrae and rib development in *Snrpb* mutants, we queried the 114 DEGs using Mammalian Phenotype (MGI) and found 12 DEGs were necessary for axial development. Pathway analysis using these 12 DEGs indicated that *Snrpb* is required for mesenchyme development (Supplementary Fig. 8a; Supplementary Data 2). Amongst these 12 DEGs we identified *Cyp26a1*, a negative regulator of the retinoic acid (RA) pathway^19,20^, as a critical candidate. RA, the active derivative of vitamin A, forms an anterior-high gradient due to its degradation in the tailbud by CYP26A1, and is crucial for somite formation and patterning^20–22^. Using RT-qPCR we showed that *Cyp26a1* expression was significantly increased in somites, paraxial mesoderm, and tailbud of *Snrpb^tmx+/–^* mutant embryos (one-tailed Student’s t-test, p < 0.05; Figure 5a). Additionally, expression of *Fgf8*, which is normally repressed by RA^23,24^, were increased in these mutant samples (one-tailed Student’s t-test, p < 0.05; Figure 5b). In contrast, we found no change in expression of *Raldh2* (Figure 5c), which encodes ALDH1A2, an enzyme that converts retinaldehyde into RA^19,20^, or the nuclear receptors (*Rara*, *Rarb*, *Rarg*, *Rxra*, *Rxrb*, and *Rxrg*) that bind to RA to regulate downstream target expression^20^ (Supplementary Fig. 8b). Together, these results suggest that elevated *Cyp26a1* expression in *Snrpb^tmx+/–^* mutant embryos perturbs the RA and FGF gradients important for somite formation and patterning.

**Figure 5.**
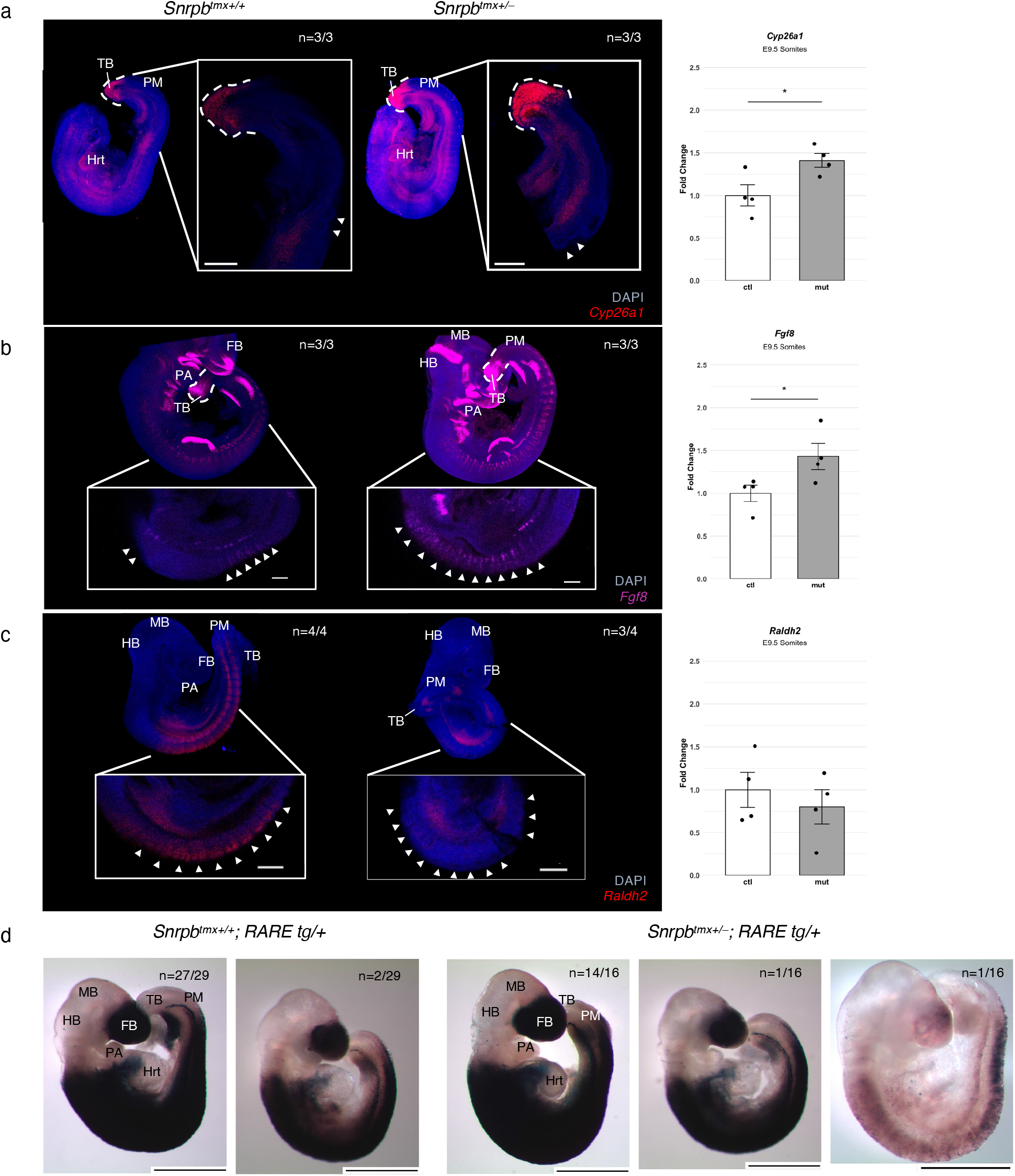
Perturbation of the retinoic acid signaling gradient in E9.5 *Snrpb^tmx+/–^* mutant somites. Representative fluorescence images of E9.5 control and *Snrpb^tmx+/–^*mutant embryos following hybridization chain reaction (HCR) using probes targeting (a) *Cyp26a1*, (b) *Fgf8*, and (c) *Raldh2*, with corresponding RT-qPCR quantification of transcript levels (mean ± SEM) in E9.5 control and mutant somites (one-tailed Student’s t-test; *p < 0.05). Arrowheads indicate somites; dashed lines delineate the tailbud region. Scale bars, 20 *µ*m. (d) Representative β-galactosidase-stained whole-mount images of E9.5 control and *Snrpb^tmx+/–^* mutant embryos carrying the RARE–LacZ reporter. FB, forebrain; MB, midbrain; HB, hindbrain; Hrt, heart; PA, pharyngeal arch; PM, presomitic mesoderm; TB, tailbud. Scale bars, 1mm.

We used hybridization chain reaction (HCR) to examine expression of *Cyp26a1*, *Fgf8*, and *Raldh2,* RA pathway transcripts important for somite development. In *Snrpb^tmx+/–^* mutant embryos, the expression domain of *Cyp26a1* was expanded in the tailbud (n=3/3; Figure 5a), while expression of *Fgf8* in somites and the tailbud was increased compared to controls (n=3/3; Figure 5b). Furthermore, *Raldh2* was not detected in somites of most mutant embryos (n=3/4, Figure 5c). To directly assess RA activity in *Snrpb^tmx+/–^* mutant embryos, we incorporated the retinoic acid response element (RARE)–LacZ reporter into our model. LacZ staining of E9.5 embryos revealed that the anterior–posterior extent of RA activity, measured as the number of somites displaying LacZ signal from the forelimb region to the tailbud, was comparable between control and mutant embryos (Supplementary Fig. 8c). However, reduced overall LacZ signal intensity was observed in a subset of *Snrpb^tmx+/–^* mutant embryos (12.5%; n=2/16) compared to a smaller proportion of controls (6.9%; n=2/29), with one mutant embryo exhibiting almost no signal (Figure 5d). Altogether, these findings suggest that while the spatial extent of RA activity is largely maintained, RA signaling is reduced in a subset of *Snrpb^tmx+/–^* mutant embryos.

### Dietary Retinoic Acid (RA) supplementation improves CCMS-like abnormalities in E12.5

#### *Snrpb^tmx+/–^* mutant embryos but exacerbates later axial defects

We hypothesized that RA supplementation during the critical window of somitogenesis would reduce the number and/or severity of axial defects. To test this, pregnant dams were fed either a control diet (Diet Gel + DMSO) or an RA-supplemented diet (Diet Gel + 100 µg/g RA in DMSO) from E7.5 to E9.5, and embryos were analyzed at E12.5 and E18.5. At E12.5, all DMSO-treated *Snrpb^tmx+/–^* mutant embryos were morphologically abnormal (n=10/10), whereas 20% of mutant embryos from RA-supplemented dams resembled control littermates (n=4/20; Supplementary Fig. 9a-c), suggesting rescue of craniofacial and limb defects normally found in mutants at this stage. However, at E18.5, all mutant embryos from RA-supplemented dams were morphologically abnormal (n=12, from 3 litters), with dome-shaped heads and smaller jaws.

Skeletal preparations revealed a significant increase in the number of axial skeletal abnormalities per mutant embryo in RA-treated compared to DMSO-treated *Snrpb^tmx+/–^* embryos (two-way ANOVA, p < 0.05; Supplementary Fig. 9d). Specifically, the proportion of *Snrpb^tmx+/–^* mutant embryos with cleft palate and discontinuous Meckel’s cartilage was higher in RA-treated mutants (n=4/9 and n=5/9, respectively) than in DMSO-treated mutants (n=1/7 for both; Supplementary Fig. 10a-b). While head size and mandible length were comparable between genotypes in the DMSO-treated group, RA-treated mutants exhibited significantly shorter mandibles (two-tailed Student’s t-test, p < 0.001; Supplementary Fig. 10c-d). Beyond the craniofacial defects, rib and vertebral abnormalities were also exacerbated in RA-treated mutants: asymmetric rib fusion was more prevalent in RA-treated mutants (n=7/9; Supplementary Fig. 11a) than DMSO-treated mutants (n=3/7; Supplementary Fig. 11b), and both abnormal ossification of thoracic vertebrae and reduced thoracic size were observed exclusively in RA-treated mutants (n=6/9; two-tailed Student’s t-test, p < 0.001; Supplementary Fig. 11c-d). Similarly, cervical vertebral defects showed higher penetrance and expressivity in RA-treated mutants than in DMSO-treated controls, including a higher proportion of embryos with missing head of T1 (n=2/9; Supplementary Fig. 12a) and pointy cervical vertebrae (n=5/9; Supplementary Fig. 12b). Altogether, these data indicate that RA supplementation during E7.5–E9.5 does not rescue CCMS-like malformations and instead exacerbates axial skeletal abnormalities in *Snrpb^tmx+/–^* mutant embryos, suggesting that *Snrpb* regulates additional developmental processes beyond RA signaling that are necessary for proper skeletal development.

### *Snrpb* loss leads to aberrant splicing of chromatin-modifying factors and altered epigenetic regulation of RA target genes in somitic tissues

As *Snrpb* encodes a core splicing factor, we used FDR < 0.05 and |ILD| > 0.05 to identify transcripts with alternative splicing events (ASEs) in control and mutant tissues. *Snrpb* mutant samples showed a small increase in the number of ASEs, with 726 splicing events in 607 genes compared to 603 splicing events in 518 genes in controls (Supplementary Fig. 13a-b; Supplementary Data 2). However, the biological processes associated with these genes showed distinct functional enrichments. Specifically, alternatively spliced genes in controls were enriched for DNA damage repair processes, whereas those in mutants were enriched for functions related to RNA modification, splicing, organelle assembly, and gene silencing, (Supplementary Fig. 13c-d). These findings indicate that at E9.5, SNRPB is required in somites, paraxial mesoderm, and/or tailbud for proper splicing of transcripts important for transcriptional regulation and post-transcriptional modifications.

We postulated that mis-splicing and/or abnormal expression of transcripts important for thorax and vertebrae development contribute to abnormalities found in *Snrpb^tmx+/–^* mutants. Therefore, we utilized the Mammalian Phenotype query on Mouse Genome Informatics (MGI) to identify genes necessary for rib and vertebrae development that were differentially spliced in *Snrpb^tmx+/–^* mutants. Using our filtering strategy, we identified a list of 52 genes with alternative splicing events. STRING analysis using the 52 genes with ASEs revealed a node that included multiple chromatin modifying proteins and significant enrichment of molecular functions such as histone binding and chromatin binding (Figure 6a; Supplementary Data 2). As *Snrpb* is required before E8.5 for proper vertebral and rib development, we reasoned that aberrant splicing of chromatin-modifying factors may disrupt epigenetic patterning and thereby underlie the skeletal defects observed in *Snrpb^tmx+/–^*mutants.

**Figure 6.**
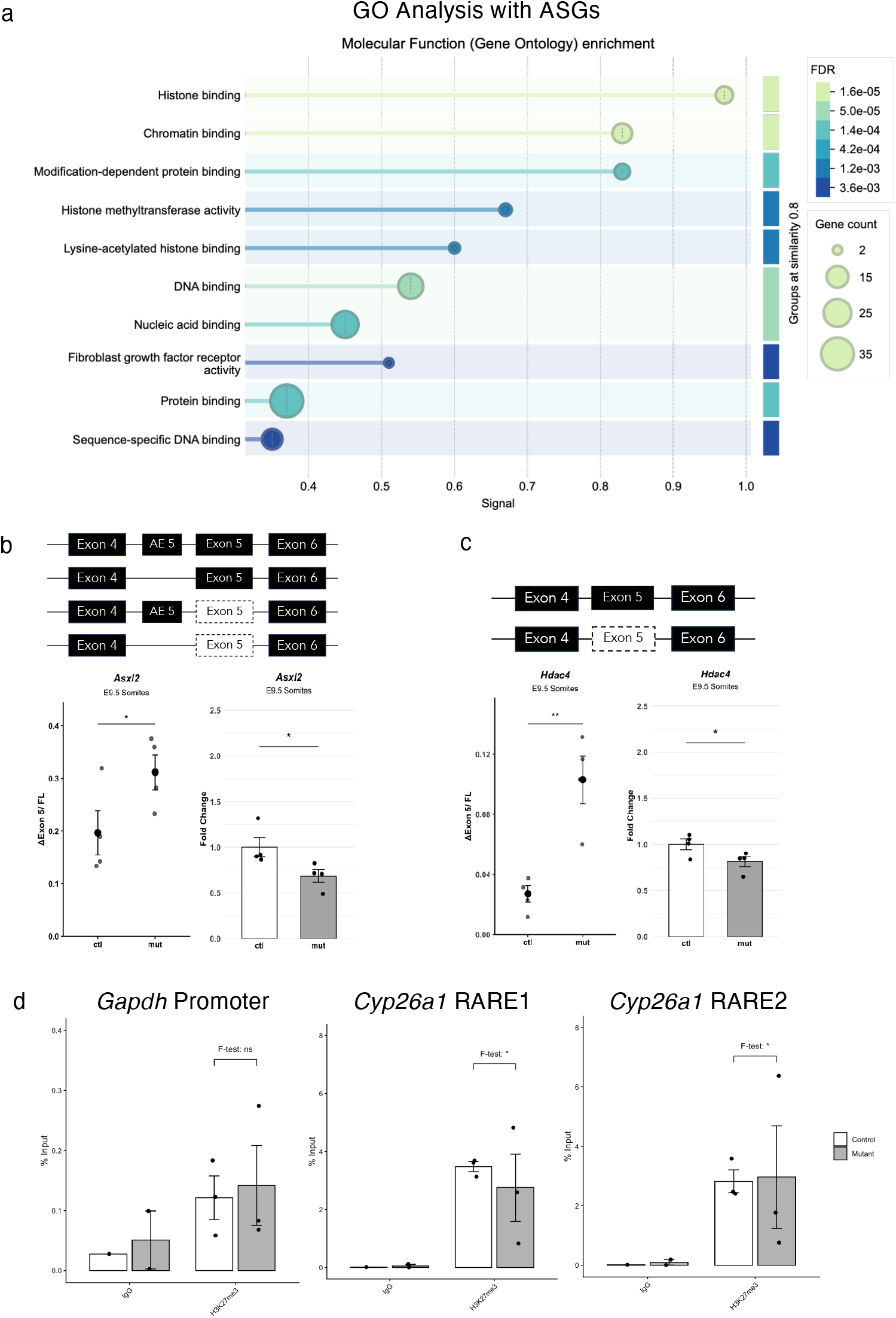
*Snrpb* loss leads to aberrant splicing of chromatin-modifying factors and altered H3K27me3 occupancy at *Cyp26a1* retinoic acid response elements. (a) Gene ontology biological process enrichment analysis of alternatively spliced genes (FDR < 0.05, |ILD| > 0.05) associated with rib and vertebral phenotypes in *Snrpb^tmx+/–^* mutants. (b) Representative RT-PCR gels and dot plots (mean ± SEM) showing increased skipping of exon 5 of *Asxl2* in E9.5 mutant somites compared to controls, and bar graphs (mean ± SEM) showing reduced *Asxl2* transcript levels in mutant samples (one-tailed Student’s t-test, *p < 0.05). (c) Representative RT-PCR gels and dot plots (mean ± SEM) showing increased skipping of exon 5 of *Hdac4* in E9.5 mutant samples compared to controls, and bar graphs (mean ± SEM) showing reduced *Hdac4* transcript levels in mutant samples (one-tailed Student’s t-test, *p < 0.05, **p < 0.01). (d) Bar graphs (mean ± SEM) showing H3K27me3 occupancy at the *Gapdh* promoter, *Cyp26a1* RARE1, and *Cyp26a1* RARE2 in E9.5 control and *Snrpb^tmx+/–^* mutant somites, paraxial mesoderm, and tailbud tissues assessed by ChIP-qPCR. Individual data points represent biological replicates. Brackets indicate one-sided F-test comparing variance between control and mutant H3K27me3 occupancy (*p < 0.05, ns = not significant).

We selected three genes encoding chromatin-modifying proteins — *Setd5*, *Asxl2*, and *Hdac4* — predicted to be alternatively spliced in *Snrpb^tmx+/–^* mutants and with known roles in vertebral differentiation and patterning, for further analysis (Supplementary Table 1). Although rMATS predicted increased inclusion of a premature termination codon (PTC)-containing alternative exon 5 of *Setd5* in mutant samples, neither exon inclusion nor transcript abundance differed significantly between control and mutant embryos by RT-PCR and RT-qPCR, respectively (Supplementary Fig. 13e). In contrast, RT-PCR revealed significantly increased skipping of constitutive exon 5 in *Asxl2*, which encodes a member of the enhancer of trithorax and polycomb protein family^25,26^, in mutants (one-tailed Student’s *t*-test, *p* < 0.05; Figure 6b). This splicing abnormality was also accompanied by reduced levels of functional *Asxl2* transcripts (one-tailed Student’s *t*-test, *p* < 0.05; Figure 6b). Similarly, RT-PCR confirmed increased skipping of constitutive exon 5 of *Hdac4*, which encodes a histone deacetylase^27^ (one-tailed Student’s *t*-test, *p* < 0.01; Figure 6c). The exon-skipping event was also associated with reduced *Hdac4* transcript levels in mutant samples (one-tailed Student’s *t*-test, *p* < 0.05; Figure 6c). Thus, we concluded that aberrant splicing of chromatin-modifying factors occurs in *Snrpb* mutant somites, presomitic mesoderm/and or tailbud, and contributes to the axial skeletal defects observed in mutant embryos.

To investigate whether the aberrant splicing of chromatin-modifying factors observed in *Snrpb^tmx+/–^* mutants alters the epigenetic landscape at RA target gene loci, we performed ChIP-qPCR using an H3K27me3 antibody on E9.5 somite, paraxial mesoderm, and tailbud samples. We first determined that H3K27me3 occupancy and variability at the *Gapdh* promoter were comparable between control and mutant embryos (Figure 6d). We next assessed H3K27me3 occupancy at two retinoic acid response elements (RAREs) associated with *Cyp26a1*: the proximal promoter RARE1, which is responsible for low-level transcriptional induction by RA, and the upstream RARE2, which acts synergistically with RARE1 to enhance the transcriptional response to RA^28^. H3K27me3 occupancy at RARE1 of *Cyp26a1* was slightly decreased in mutant tissues compared to controls, and occupancy at both RARE sites showed greater variability in mutants compared to controls (one-tailed F-tests, *p* < 0.05; Figure 6d). These findings suggest that aberrant splicing of chromatin-modifying factors in *Snrpb^tmx+/–^* mutants alters the epigenetic regulation of *Cyp26a1* and may contribute to its elevated expression through reduced repressive H3K27me3 marks at RA response elements.

## Discussion

While previous studies have provided insight into how mutations in *SNRPB* impair osteogenesis and craniofacial development, this study establishes the first mouse model that recapitulates the broader spectrum of axial skeletal abnormalities associated with Cerebro-Costo-Mandibular Syndrome (CCMS), including cleft palate, micrognathia, a bell-shaped thorax, and posterior rib gaps. By inducing heterozygous deletion of *Snrpb* at distinct embryonic stages, we show that SNRPB levels are required between E8.0-E8.5 for axial skeletal development. Nevertheless, phenotypic variability is an inherent feature of this model. Mice mate at variable times during the night, and embryos collected at specific timepoints after tamoxifen injection spanned a range of approximately 10 somites across three litters, reflecting developmental stage differences at the time of *Snrpb* deletion both between and within litters. Given that SNRPB is required within a narrow developmental window, even small differences in the stage at which *Snrpb* deletion is induced are likely to contribute to variability in the penetrance and severity of axial skeletal defects, as well as variability in gene expression and alternative splicing patterns observed across mutant embryos.

Despite this variability, *Snrpb^tmx+/–^* mutant embryos consistently formed somites normally and expressed *Pax1*, *Uncx4.1*, and *Tbx18* in their appropriate domains at E9.5, yet developed rib and vertebral defects. This dissociation suggests information that determines the position and identity of axial skeletal elements is established well before those elements form, and that its disruption is not reported by the markers conventionally used to assess sclerotome specification and rostrocaudal somite polarity. These findings highlight the importance of SNRPB-mediated splicing during early somitogenesis and suggest that axial skeletal defects in CCMS arise from disruption of early developmental programs.

To identify molecular changes that precede overt morphological abnormalities, we performed bulk RNA-seq on E9.5 somites, paraxial mesoderm, and tailbud tissues, a stage at which mutant embryos appear morphologically normal but have undergone Cre-induced deletion. We propose that disruption in multiple pathways likely contributes to the axial skeletal abnormalities in mutants: p53 pathway activation induced cell death, dysregulation of retinoic acid (RA) signaling, and alternative splicing of chromatin modifiers (Figure 7).

**Figure 7.**
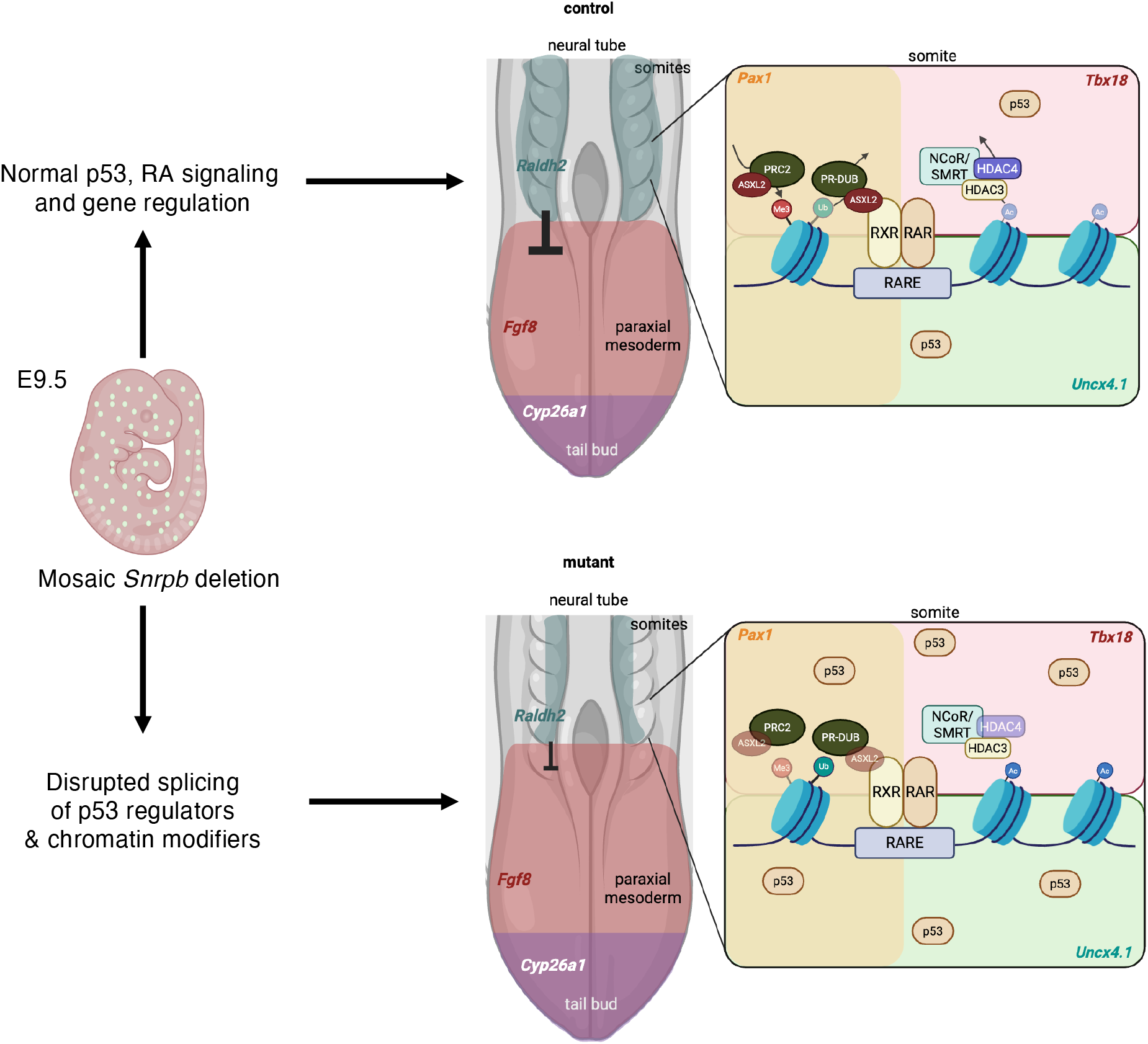
Molecular consequences of mosaic *Snrpb* deletion in E9.5 somites. Schematic comparing molecular states in control and *Snrpb^tmx+/–^*mutant embryos at E9.5. In controls (top), *Raldh2* expression in somites and paraxial mesoderm establishes an RA gradient that, together with CYP26A1-mediated degradation in the tailbud, restricts *Fgf8* expression in the tailbud. Within somites, ASXL2 functions in both the PRC2 and PR-DUB complexes to maintain H3K27me3 (Me3) and regulate H2AK119ub1 (Ub) levels at target loci, while HDAC4 scaffolds the NCoR/SMRT–HDAC3 complex to repress gene expression via histone deacetylation. p53 levels are low, *Pax1* marks the sclerotome, and *Tbx18* and *Uncx4.1* expression demarcate normal rostrocaudal somite polarity. In *Snrpb^tmx+/–^* mutants with mosaic *Snrpb* deletion at E9.5 (bottom), aberrant splicing of *Asxl2* and *Hdac4* disrupts chromatin regulation, leading to reduced H3K27me3 occupancy at retinoic acid response elements (RAREs) and impaired NCoR/SMRT–HDAC3 recruitment, likely resulting in increased histone acetylation (Ac). Reduced *Raldh2* expression in somites and increased *Cyp26a1* and *Fgf8* expression in the tailbud disrupt the RA and FGF gradients important for somite patterning. Elevated nuclear p53 in anterior somites reflects increased cellular stress that later results in increased cell death. *Pax1*, *Tbx18*, and *Uncx4.1* expression remain unchanged, indicating that sclerotome specification and rostrocaudal somite polarity are maintained despite these molecular disruptions. RXR, retinoid X receptor; RAR, retinoic acid receptor; RARE, retinoic acid response element; Me3, H3K27me3; Ub, H2AK119ub1; Ac, acetylation.

Consistent with previous findings in *Snrpb*-deficient neural crest cells^10^, we observed altered splicing of the p53 negative regulator *Mdm2*, increased expression of a p53 target gene *Phlda3*, and a significantly greater proportion of p53-positive cells in anterior than posterior somites of *Snrpb^tmx+/–^* mutants, suggesting that p53 pathway activation is region-specific and may contribute to the more severe axial skeletal defects observed in anterior ribs and vertebrae relative to posterior regions. However, this early p53 activation was not immediately associated with cell death in E9.5 anterior somites. Instead, cell death was detected later, in E12.5 anterior somite-derived tissues. Because the magnitude and duration of p53 activation are known to influence cellular outcomes, with transient activation promoting stress adaptation or cell-cycle arrest and sustained activation triggering apoptosis, early p53 activation in mutants may reflect a transient stress response that precedes later cell death^29^. Thus, p53-associated stress responses may contribute to axial abnormalities, but they are unlikely to fully explain the penetrance or severity of these defects.

In addition to p53 pathway activation, we identified alternative splicing of several chromatin modifiers, including *Asxl2* and *Hdac4*, both of which play important roles in transcriptional regulation during axial skeletal development. ASXL2 is a core component of the Polycomb Repressive Deubiquitinase (PR-DUB) complex and Polycomb Repressive Complex 2 (PRC2)^30^. The PR-DUB complex removes PRC1-deposited H2AK119ub1 marks to fine-tune Polycomb-mediated gene repression. As PRC1 and PRC2 act cooperatively to maintain repressive chromatin domains: PRC1-mediated H2AK119ub1 can recruit PRC2 to deposit H3K27me3. Loss of ASXL2 would be expected to compromise PR-DUB and PRC2 activity, leading to accumulation of H2AK119ub1, disruption of PRC1-PRC2 balance, and altered H3K27me3 occupancy at Polycomb target genes critical for axial skeletal patterning^31–35^. In contrast, reduced HDAC4 likely impairs NCoR/SMRT–HDAC corepressor activity, leading to increased histone acetylation^36–38^. Together, these defects are predicted to destabilize repressive chromatin states and contribute to widespread transcriptional dysregulation during axial skeletal development.

The alternative splicing of chromatin modifiers is particularly relevant in the context of the RA signaling defects observed in E9.5 mutants. RA signaling is tightly linked to chromatin regulation, as RA receptors regulate transcription through the recruitment of co-activator and co-repressor complexes that include histone acetyltransferases and histone deacetylases. Therefore, perturbation of chromatin regulators such as HDAC4 could alter the ability of RA receptors to properly regulate target genes^20^. In addition, the ASXM2 domain of ASXL2 mediates interaction with RXRα^39^, suggesting that ASXL2 may influence nuclear receptor-dependent transcriptional programs, including those involved in retinoid signaling. Consistent with this, we observed altered H3K27me3 occupancy and increased variability at the RARE1 and RARE2 sites of *Cyp26a1* in *Snrpb^tmx+/–^* mutant somites, suggesting that aberrant splicing of chromatin-modifying factors may directly alter the epigenetic regulation of RA target genes. Reduced H3K27me3 at RARE1 would be expected to relieve repression of *Cyp26a1*, consistent with the elevated *Cyp26a1* expression observed in mutant somites and the consequent perturbation of the RA gradient. Conversely, broader disruption of chromatin regulation could also directly affect transcription of genes involved in RA synthesis, degradation, or downstream signaling. Together, these findings raise the possibility that splicing defects in chromatin modifiers and altered RA signaling cooperate to disrupt transcriptional programs required for axial skeletal patterning, with epigenetic dysregulation at RA response elements representing one potential mechanistic link between these two pathways.

This interaction between chromatin dysregulation and RA signaling may help explain why maternal RA supplementation alone failed to rescue the axial skeletal phenotypes in mutants. If RA pathway disruption is only one component of a broader transcriptional defect, restoring RA levels may be insufficient to normalize downstream gene expression. In addition, systemic maternal RA supplementation does not selectively increase RA activity in mutant cells. Because tamoxifen-induced recombination is mosaic, not all cells are expected to undergo *Snrpb* deletion or become RA deficient. As a result, dietary RA supplementation may increase RA signaling in both mutant and non-mutant neighboring cells, potentially creating local regions of excess RA activity. Since both RA deficiency and excess RA are known to disrupt embryonic patterning^40^, this lack of cell-type specificity may contribute to the failure of RA supplementation to rescue the phenotype and may explain why axial defects were exacerbated at later developmental stages.

Unexpectedly, supplementation of Diet Gel with DMSO alone partially alleviated skeletal phenotypes in mutants, resulting in improved embryo weights and reduced severity of axial abnormalities. Although the mechanism underlying this effect remains unclear, DMSO has been shown to influence multiple cellular processes even at low concentrations. In zebrafish, DMSO exposure can alter developmental morphology and physiology, including processes involved in somitogenesis^41^. In mammalian systems, DMSO has been reported to influence chromatin structure by increasing histone acetylation and altering gene expression, thereby perturbing developmental transcriptional programs^42,43^; indeed, DMSO alone is sufficient to shift chromatin state and drive trophectoderm differentiation and blastoid formation from human naïve pluripotent stem cells^44^. DMSO has also been shown to suppress inflammatory and stress-associated signaling pathways^45^. As a result, partial phenotypic improvement observed in our mutants may reflect DMSO-mediated modulation of stress response, chromatin state, or transcriptional regulation. Future studies examining the effects of DMSO on gene expression, chromatin accessibility, and cellular stress during early embryogenesis will be necessary to determine how DMSO modifies developmental outcomes in this model.

In summary, our study demonstrates that reduced SNRPB disrupts early developmental processes through combined effects on cellular stress responses, RNA splicing, chromatin regulation, and RA signaling. These perturbations arise at a stage when somites form and are patterned normally and manifest only later as the vertebral and rib malformations characteristic of CCMS. Our findings therefore suggest that the developmental information specifying axial skeletal identity is laid down early in the somite and is vulnerable to spliceosomal dysfunction in ways that conventional markers of sclerotome specification and rostrocaudal somite polarity do not reveal. This physiologically relevant *in vivo* model of CCMS provides a foundation for identifying the gene regulatory programs through which SNRPB acts.

## Materials and Methods

### Mouse Lines

All procedures and experiments were performed according to the guidelines of the Canadian Council on Animal Care and approved by the Animal Care Committee of the McGill University Health Centre Research Institute. Generation of the conditional *Snrpb* mouse line with loxP sequences flanking exons 2 and 3 has been previously described^10^. The *Snrpb* conditional knockout strain is maintained on a mixed C57BL/6J and CD1 genetic background. The *mT/mG* [*Gt(ROSA)26Sort^m4(ACTB-tdTomato,-EGFP)Luo^*/J] (strain#: 007676) was purchased from The Jackson Laboratory. *R26-CreERT2* [*B6.129-Gt(ROSA)26Sor^tm1(cre/ERT2)Tyj^/J*] (strain#: 008463) was a kind gift from Dr. Hugh Clarke (Department of Obstetrics and Gynecology, McGill University, Montreal, Canada). *RARE-hsp68LacZ* [*Tg(RARE-Hspa1b/lacZ)12Jrt/J*] (strain#: 008477) was a kind gift from Dr. Celeste Nelson (Department of Chemical & Biological Engineering, Princeton University, NJ, USA).

### Collection and Genotyping of Embryos

Females were placed with a male overnight and checked for vaginal plug in the morning. Noon on the day of the plug was considered embryonic day (E) 0.5. Dissections were carried out under a Leica stereomicroscope (Leica MZ6). On day of collection, the embryos were removed from their extraembryonic membranes, and the yolk sac was collected for DNA extraction and genotyping. For embryos collected from E8.5 to E10.5, the pairs of somites were counted. Embryos were collected in 1×PBS, fixed in 4% PFA or 1% PFA and stored in 1×PBS at 4 °C unless otherwise specified.

### Tamoxifen Treatment

Pregnant females received a single intraperitoneal injection of tamoxifen (Sigma T5648) and progesterone (Sigma P8783) at doses of 2 mg and 1 mg per 20 g body weight, respectively. Progesterone was co-administrated to reduce tamoxifen-induced miscarriage as previously described^46,47^. Tamoxifen (10 mg/mL) and progesterone (5 mg/mL) were prepared together in sterile corn oil (Sigma C8267), sonicated for 30 minutes to ensure complete dissolution, protected from light, and stored at 4°C for up to one week.

### Retinoic Acid (RA) Dietary Supplementation

For retinoic acid (RA) supplementation using 76A DietGel (ClearH2O DietGel® 76A), RA was first dissolved in DMSO (Thermo Scientific, J66650.AD) to generate a 10 mM stock solution, then incorporated into DietGel at a final concentration of 100 µg/g (6.66 µL per 20 g DietGel). For vehicle controls, DMSO was added at the same volume (6.66 µL per 20 g; 0.033% v/w). DietGel, vehicle-, or RA-supplemented DietGel replaced standard chow from E7.5–E9.5.

### Cartilage and Skeletal Preparation of Embryos

To investigate cartilage formation, E14.5 embryos were stained with Alcian Blue. For skeletal staining, the skin was removed from frozen E17.5/E18.5 embryos and stained as previously described^48^. Cartilage and skeletal preparations were imaged and analyzed using a Leica stereomicroscope (Leica MZ6). Measurements were performed using Fiji (ImageJ), and statistical analyses were conducted in R (RStudio).

### Embryo Embedding and Sectioning

For cryo-embedding, fixed embryos were cryoprotected in 30% sucrose overnight, embedded in cryomatrix, and sectioned sagittally at 10 μm thickness. For embedding in paraffin, fixed embryos were washed in 1xPBS then dehydrated to 100% EtOH and embedded using paraffin. Embedded embryos were sectioned at 10 μm thickness on a Leica RM2155 microtome and mounted on positively charged slides for further analysis.

### TUNEL Assay

Cryosections and paraffin sections of embryos fixed overnight in 4% PFA were used for TUNEL assay. TUNEL assay was carried out using a Cell Death Detection Kit, TMR Red (Roche 12156792910) as per the manufacturer’s protocol. Slides were mounted with Fluoroshield™ with DAPI (Sigma F6057) to visualize the nuclei. Images were captured on a Leica microsystem (model DM6000B) and Leica camera (model DFC 450). For quantification of TUNEL signal, particle analysis on Image J was used. Anterior somites denote those found rostral to the forelimb limb buds whereas somites posterior somites refer to those found caudal to the forelimb limb buds.

### Immunohistochemistry (IHC)

Cryosections sections of embryos fixed overnight in 1% PFA were used for IHC. Embryos were sectioned at 10-μm thickness for immunohistochemistry as previously described^49^. Anti-P53 primary antibody (2524T, Cell Signaling Technology; 1:250 dilution) or Anti-PHH3 primary antibody (06-570, Millipore Sigma; 1:200 dilution) was used. A VECTASTAIN® Universal Quick HRP Kit was used as secondary antibody and visualized with diaminobenzidine (DAB; AB64238, Abcam). Images were captured on a Leica microsystem (model DM6000B) and Leica camera (model DFC 450). For quantification of IHC signal, particle analysis on Image J was used. Anterior and posterios somites were annotated as described above.

### Hybridization Chain Reaction (HCR)

Hybridization Chain Reaction (HCR) was performed using the v3.0 or Gold RNA-FISH kit (Molecular Instruments) according to the manufacturer’s instructions. Images were acquired using a Zeiss LSM780 laser scanning confocal microscope equipped with IR lasers and an optical parametric oscillator (OPO) at the Molecular Imaging Platform of the Research Institute of the McGill University Health Centre (RI-MUHC). Image analysis was performed using ZEISS ZEN Lite software.

### Whole Mount *in situ* Hybridization (ISH)

Fixed embryos were dehydrated using a graded methanol series for wholemounts. Wholemount RNA *in situ* hybridization was performed as previously described^50^.

### Wholemount X-Gal Staining

Embryos carrying *RARE-hsp68LacZ* were stained with freshly prepared X-gal staining solution overnight at 37°C in the dark as previously described^49^. Post-staining, embryos were embedded in Cryomatrix and stored at −80°C until sectioning.

### RNA Isolation

To isolate somites and tail bud for RNA, the head, limb buds, and heart were removed from 25– 28 somite-stage E9.5 embryos. All post-otic somites, presomitic mesoderm, and tail bud were collected in 250 μl RNAlater (Invitrogen AM7020). Somites from four embryos were pooled for RNA extraction using the Qiagen RNeasy micro kit per the manufacturer’s instructions (Qiagen 74104). Three pools each of wild-type and mutant somites were used for RNA sequencing. For E12.5 samples, the head, heart, and limb buds were removed and trunk tissues were flash-frozen on dry ice. Trunks from two embryos were pooled per sample, and RNA was extracted using TRIzol reagent according to the manufacturer’s instructions (Invitrogen 15596026).

### RNA Sequencing and Analysis

Sequencing libraries were prepared by the McGill Genome Centre (Montréal, Canada) using the TruSeq Stranded Total RNA Sample Preparation Kit (TS-122-2301; Illumina, San Diego, CA, USA), comprising ribosomal RNA depletion, RNA fragmentation, first-and second-strand complementary DNA (cDNA) synthesis, 3′-end adenylation, adaptor ligation, and PCR enrichment of adaptor-ligated fragments. Libraries were sequenced on an Illumina NovaSeq 6000 instrument in paired-end mode (PE100; 2 × 100 nt), yielding between 109 and 230 million paired-end reads per sample. Raw reads were adaptor-and quality-trimmed with Cutadapt^51^ and aligned to the mouse reference genome (GRCm38/mm10) with the STAR aligner^52^ (v2.6.1d) under default parameters, guided by the GENCODE^53^ M2 annotation (release M2, 2013). Gene-level expression was quantified with htseq-count from the HTSeq framework^54^ (v0.13.5).

Differential splicing analysis was performed with rMATS^55^ (v4.1.2). Candidate events were filtered by excluding those with a mean inclusion-junction count below 5 in either the wild-type (WT) or heterozygous samples. A differentially spliced event (DSE) was defined by an absolute inclusion-level difference (|ΔΨ|) greater than 0.05 together with a Benjamini-Hochberg false discovery rate (FDR) below 0.05. This comparatively permissive FDR threshold was applied deliberately to obtain a dataset broadly enriched for alternative-splicing events, thereby enabling the detection of global tendencies - such as an increased propensity for exon skipping or intron retention in the mutants. To characterise 3′ splice-site sequences, branchpoints (BPs) located upstream of DSEs were predicted with LaBranchoR^56^, a deep-learning tool that employs a bidirectional long short-term memory (LSTM) network for BP prediction. Consensus motifs spanning the predicted BPs and their flanking regions were generated with WebLogo^57^ (v3.0).

Differential expression analysis (DEA) was conducted with the DESeq2^58^ package. Significant differentially expressed genes (DEGs) were defined by an FDR below 0.05, and a fold change threshold of 1 imposed on the absolute fold change (FC) so as to retain even subtle expression changes. For Kyoto Encyclopedia of Genes and Genomes (KEGG)^59^ pathway enrichment, the combined set of up-and down-regulated DEGs was supplied as the query to the g:Profiler^60^ framework as implemented in the gprofiler2^61^ R package (gost function), with all genes detected in the DEA used as the statistical background.

DEGs (FDR-adjusted *P* < 0.05; fold change > 1) and alternatively spliced genes (ASGs; FDR < 0.05, |ΔΨ| > 0.05) were analysed as separate datasets with the Mouse Genome Informatics (MGI)^62^ Batch Query tool. For each dataset, genes annotated with rib and vertebral abnormalities under the Mammalian Phenotype (MP) Ontology^63^ were identified and extracted. The resulting DEG-and ASG-derived gene sets were then analysed independently in STRING^64^ to reconstruct protein–protein interaction networks.

### Quantitative RT-PCR

Total RNA was treated with DNAse (NEB; according to the manufacturer’s protocol) and used for reverse transcription with iScript™ Reverse Transcription Supermix for RT. Quantitative RT-PCR was performed using Advanced Universal SYBR® Green Supermix (Bio-Rad 1725271). Experiments were performed in duplicates to ensure technical replicability. Target genes were normalized with the normalization factor as calculated by geNorm software^65^. Three housekeeping genes – *Actb*, *Gapdh,* and *Sdha* – were used for generation of the normalization factor.

### Primers

See Supplementary Table 2 for specific primer sequences. Primers for alternative splicing events in *Mdm2* and *Mdm4*, as well as qPCR primers for *Sdha*, *Trp53inp1*, *Ccng1*, and *Phlda3*, were previously described^49^.

### ChIP-qPCR

For ChIP-qPCR assays, E9.5 trunk tissues (head and heart removed) were pooled from three wild-type and three *Snrpb^tmx+/–^* mutant embryos (25–28 somites) per biological replicate. Tissues were homogenized in cold PBS containing 10 mM EDTA, 0.05% Triton X-100, and PMSF, then crosslinked in 1% formaldehyde in PBS for 10 minutes at room temperature and quenched with 125 mM glycine. Chromatin was isolated through sequential lysis in hypotonic buffer (0.25% Triton X-100, 10 mM Tris pH 8.0, 10 mM EDTA, 0.5 mM EGTA, protease inhibitor cocktail) and high-salt buffer (200 mM NaCl, 10 mM Tris pH 8.0, 1 mM EDTA, 0.5 mM EGTA, protease inhibitor cocktail), then sonicated in SDS lysis buffer (0.5% SDS, 0.5% Triton X-100, 10 mM Tris pH 8.0, 150 mM NaCl, 1 mM EDTA, 0.5 mM EGTA, protease inhibitor cocktail) using a probe sonicator (10 s on, 30 s off, 3 cycles) to yield fragments of 200–700 bp, as verified by agarose gel electrophoresis. Chromatin was diluted 1:5 in dilution buffer (0.5% Triton X-100, 10 mM Tris pH 8.0, 150 mM NaCl, 2 mM EDTA) and immunoprecipitated overnight at 4°C using protein A/G beads pre-incubated with 1 µg of anti-H3K27me3 antibody (Invitrogen, MA5-11198) or control IgG (Cell Signaling, 2729S). Following sequential washes, crosslinks were reversed overnight at 65°C, and DNA was purified using a PCR purification kit (QIAGEN) and eluted in 30 µL DEPC-treated water. Quantitative PCR was performed using primers targeting the RARE1 and RARE2 sites of *Cyp26a1* (Supplementary Table 2). Results were normalized to input chromatin. ChIP protocol was adapted from Jabado laboratory, McGill University.

## Acknowledgements

We would like to thank the McGill Genome Centre for running the RNA-seq. We thank Cassandra Millet-Boureima for her experimental contributions to this study, and members of the Majewska laboratory for their helpful comments on the manuscript. We thank the Fonds de recherche du Québec – Santé (336114) and the Research Institute of McGill University Health Centre for supporting Y.D. We would also like to acknowledge the professional and technical support from the Animal Resource Division of Research Institute -McGill University Health Centre for maintaining our mouse colonies. L.A.J-M. and J.M. are members of the Research Institute of the McGill University Health Centre, which is funded in part by Fonds de Recherche du Quebec en Santé (FRQS).

## Author contributions

L.A.J.-M. designed research; Y.D., M.P., J.S., and S.S.A. performed research; Y.D., E.B., and J.M. analyzed data; and Y.D. and L.A.J.-M. wrote the paper.

