## Supplementary Figures for "Developmental pleiotropy revealed by mosaic heterozygous *Snrpb* deletion underlies CCMS-like axial skeletal defects"

### Supplementary Figure 1.

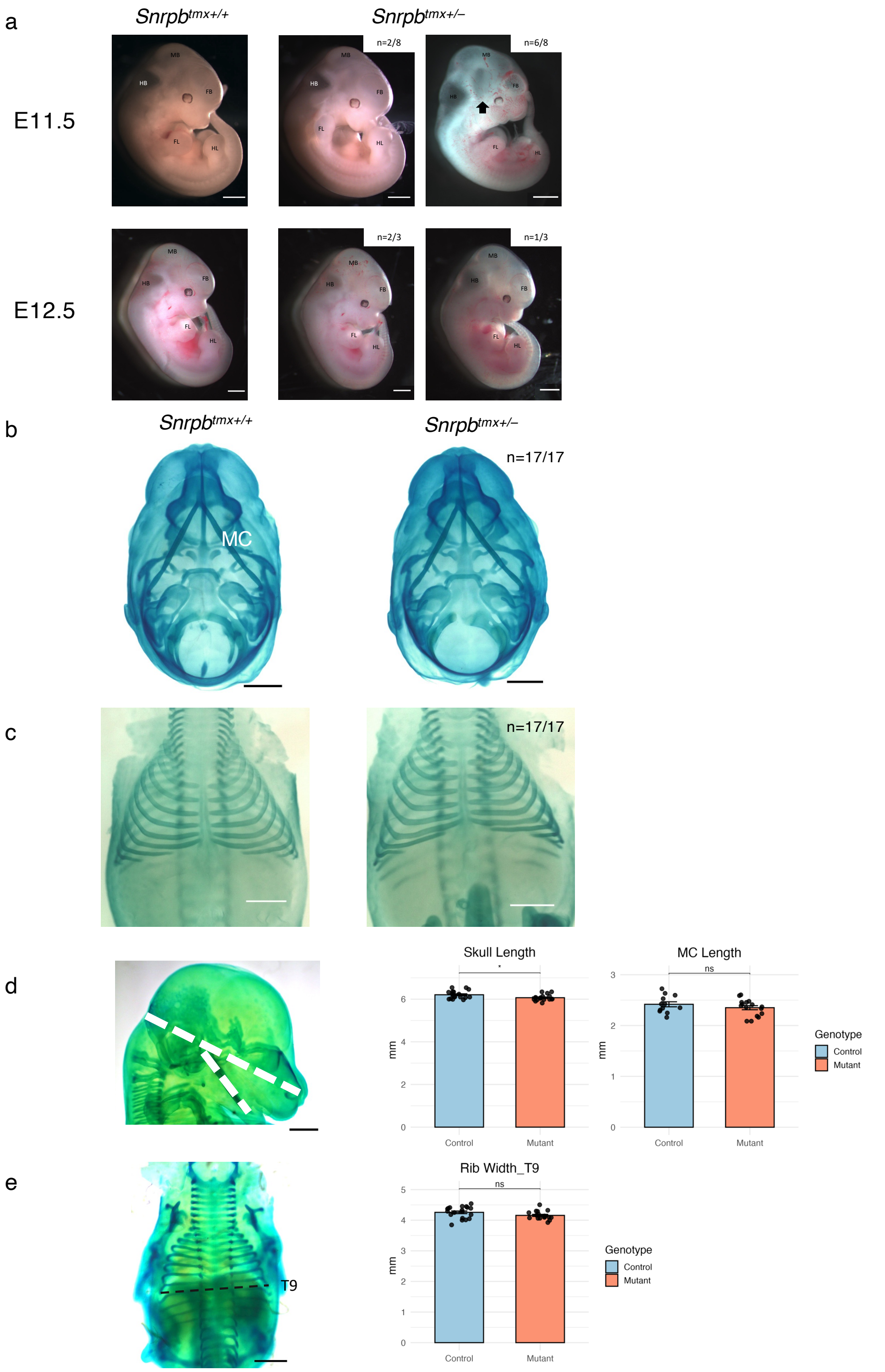


**Supplementary Figure 1. Phenotypic outcomes of tamoxifen-induced *Snrpb* deletion at E8.0 and after E8.5.**

(a) Representative images of E11.5 and E12.5 control and *Snrpb*$^{tmx+/-}$ mutant embryos with tamoxifen-induced *Snrpb* deletion at E8.0. At E11.5, 6 out of 8 mutant embryos exhibited reduced forebrain, expanded cephalic flexure (arrowheads), and reduced forelimb and hindlimb development. At E12.5, 2 out of 3 mutants displayed reduced forebrain; one mutant appeared smaller overall with reduced forebrain. FB, forebrain; MB, midbrain; HB, hindbrain; FL, forelimb; HL, hindlimb. Scale bars, 1mm. Representative cartilage preparations of E14.5 control and *Snrpb*$^{tmx+/-}$ mutant embryos with tamoxifen-induced *Snrpb* deletion between E8.5 and E9.5. Mutant embryos (n=14/14) exhibit (b) craniofacial and (c) rib patterning comparable to controls. MC, Meckel's cartilage. Scale bars, 1mm. Bar plots (mean ± SEM) comparing (d) skull length, Meckel's cartilage length, and (e) T9 rib width between control and mutant embryos (two-tailed Student's t-test, *$p < 0.05$).

**Supplementary Figure 2.**

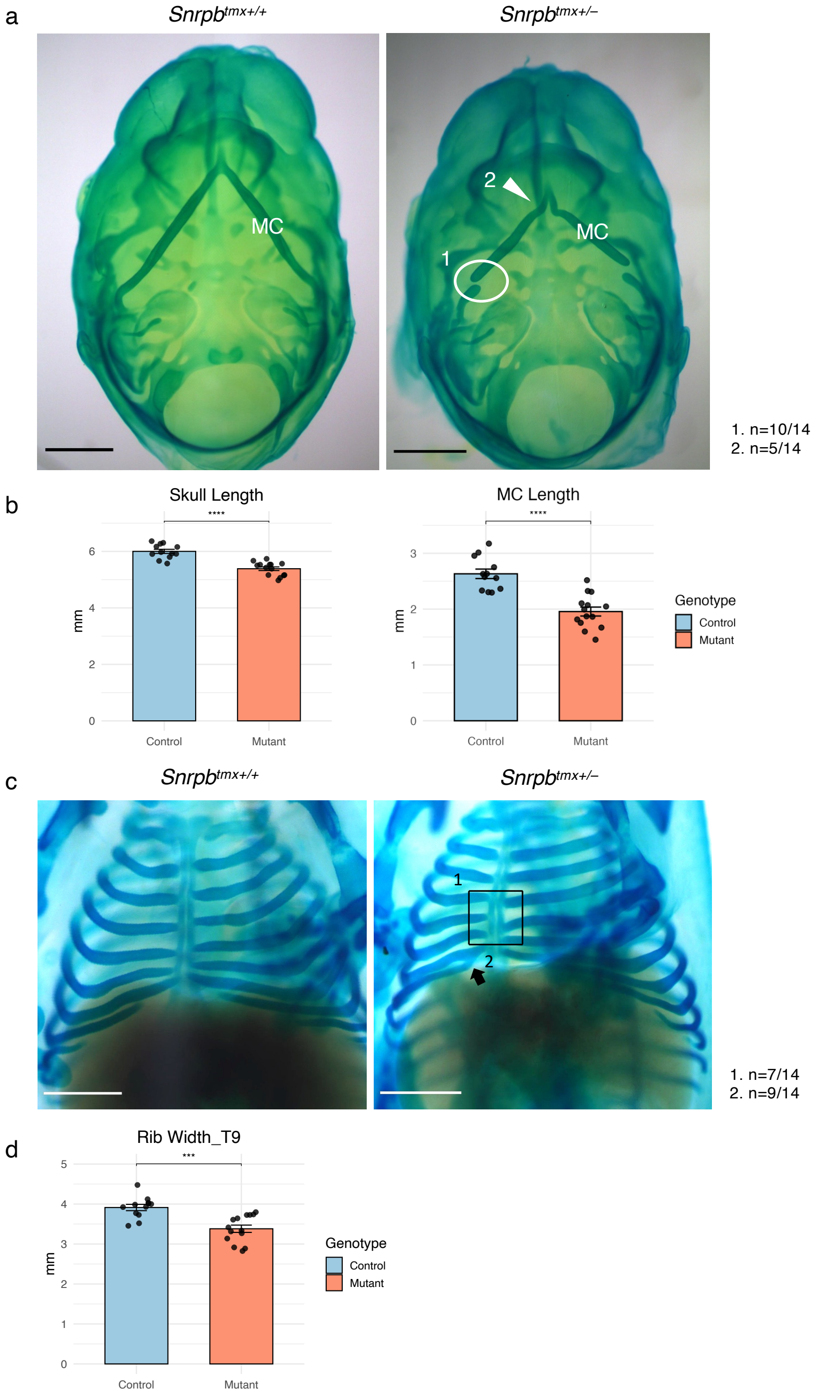


**Supplementary Figure 2. Craniofacial and rib abnormalities in E14.5 *Snrpb*$^{tmx+/-}$ mutants.**
(a) Representative cartilage preparations showing lower jaw abnormalities in *Snrpb*$^{tmx+/-}$ mutant embryos, including 1) disconnection of Meckel's cartilage from the middle ear structures (n=10/14) and 2) curvature of Meckel's cartilage (n=5/14). (b) Bar plots (mean ± SEM) comparing skull length and Meckel's cartilage length between control and mutant embryos (two-tailed Student's t-test, ****$p < 0.0001$). (c) Representative cartilage preparations showing rib abnormalities in *Snrpb*$^{tmx+/-}$ mutant embryos, including 1) asymmetric rib fusion to the sternum (n=7/14) and 2) false rib fusions (n=9/14). (d) Bar plot (mean ± SEM) comparing T9 rib width between control and mutant embryos (two-tailed Student's t-test, ***$p < 0.001$). Scale bars, 1mm.

### Supplementary Figure 3.

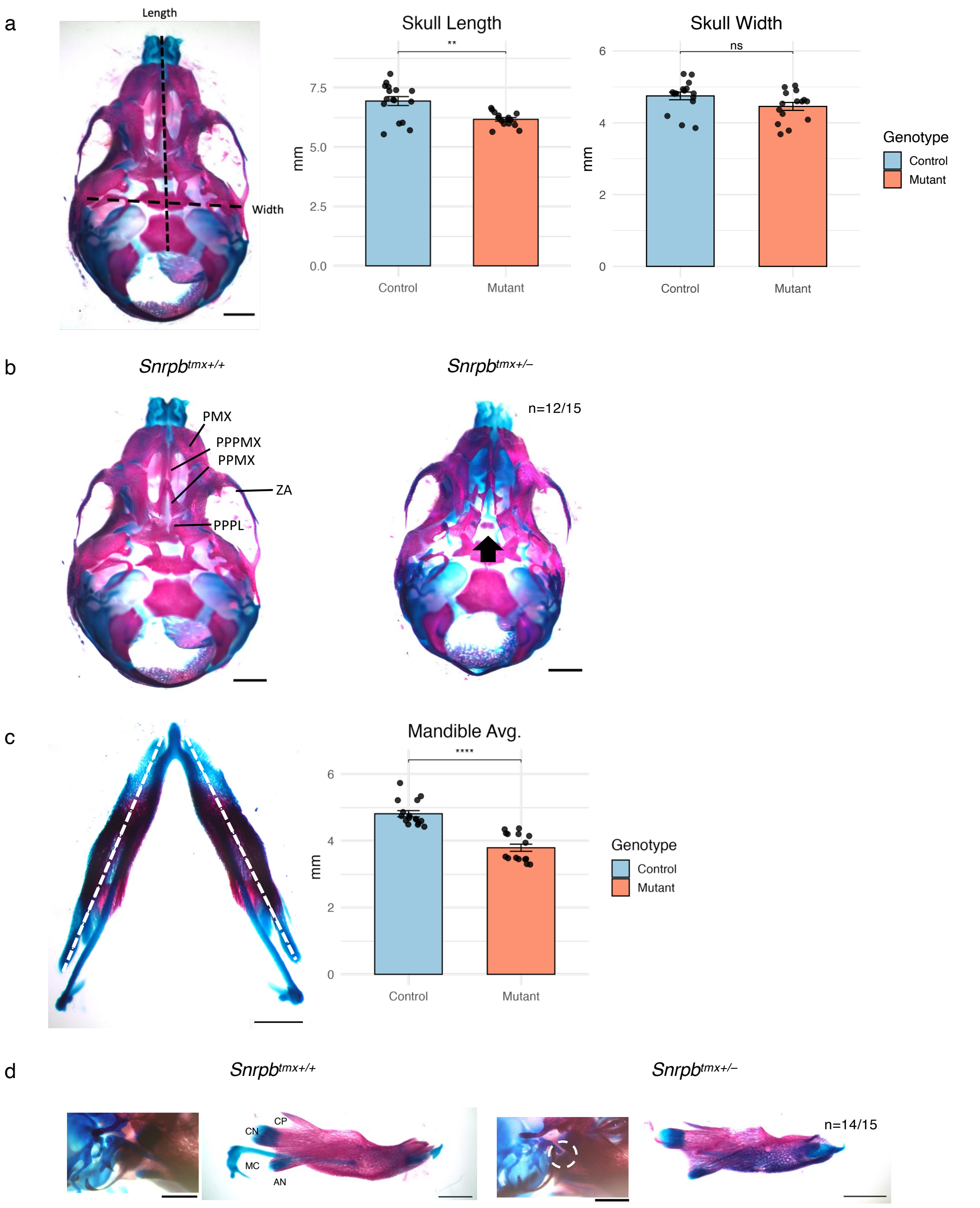


**Supplementary Figure 3. Craniofacial abnormalities in E17.5 *Snrpb*$^{tmx+/–}$ mutants.**
(a) Bar plots (mean ± SEM) comparing skull length and width between control and *Snrpb*$^{tmx+/–}$ mutant embryos (two-tailed Student's t-test, **$p < 0.01$). (b) Representative ventral skeletal preparations of control and *Snrpb*$^{tmx+/–}$ mutant skulls showing cleft palate (arrowheads; n=12/15). PMX, premaxilla; PPMX, palatal process of maxilla; PPPMX, palatal process of premaxilla; ZA, zygomatic arch; PPPL, palatal process of palatine. (c) Bar plot (mean ± SEM) comparing averaged mandible length between control and *Snrpb*$^{tmx+/–}$ mutant embryos (two-tailed Student's t-test, ***$p < 0.001$). (d) Representative skeletal preparations of control and *Snrpb*$^{tmx+/–}$ mutant embryos showing Meckel's cartilage disconnected from the middle ear structures (circles; n=14/15) and shortened mandibles. CP, coronoid process; CN, condylar process; AN, angular process; MC, Meckel's cartilage. Scale bars, 1mm.

**Supplementary Figure 4.**

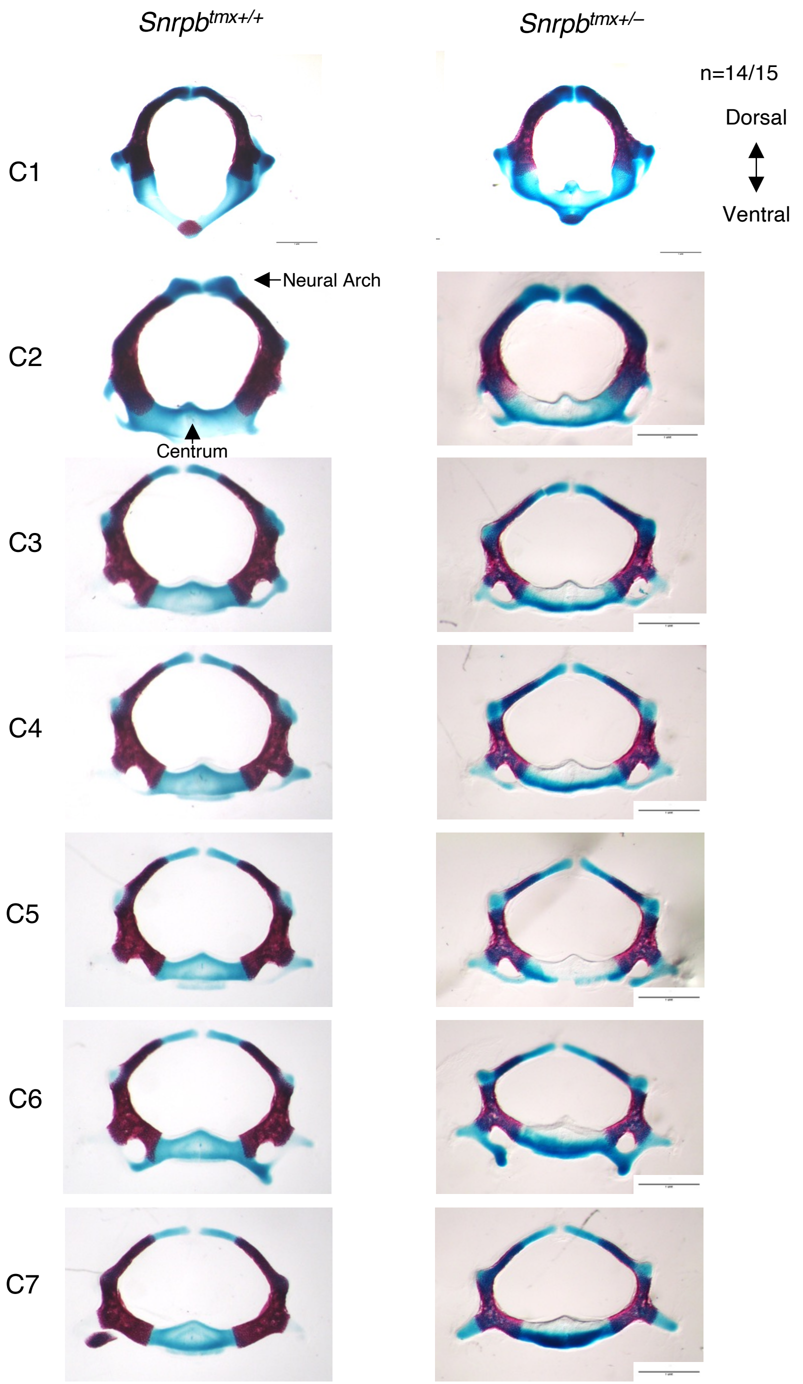


**Supplementary Figure 4. Cervical vertebral abnormalities in E17.5 *Snrpb*$^{tmx+/–}$ mutants.** Representative skeletal preparations of cervical vertebrae from control and *Snrpb*$^{tmx+/–}$ mutant embryos. In 14 out of 15 mutant embryos, the C1 vertebra exhibited reduced dorsoventral height, while the C3–C7 vertebrae displayed elongated centra and more tapered neural arches. Scale bars, 1mm.

**Supplementary Figure 5.**

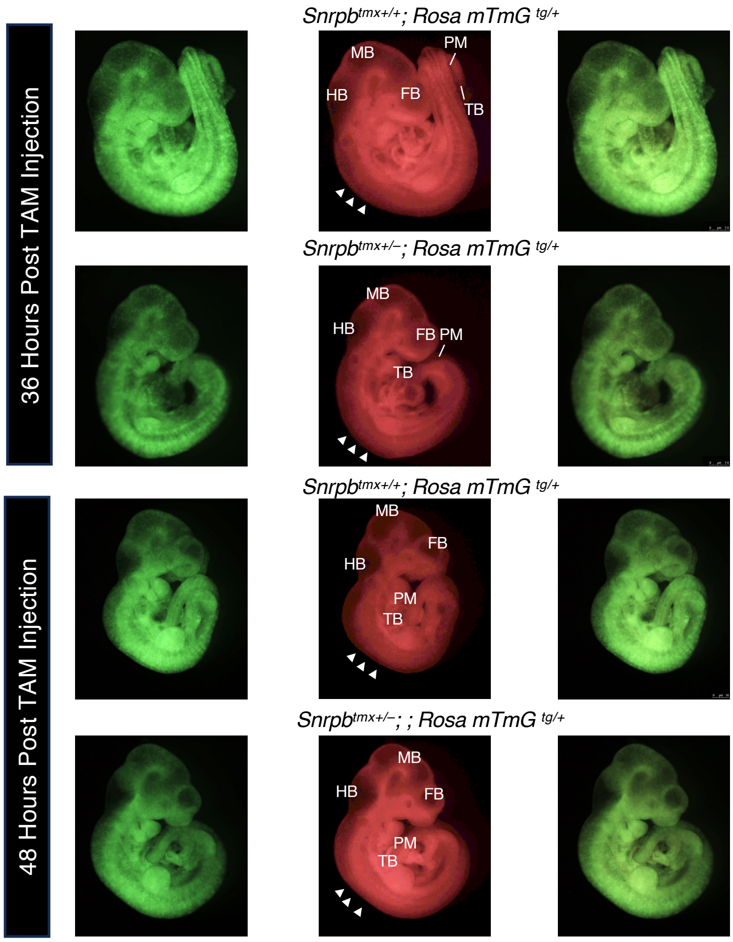


**Supplementary Figure 5. Persistent mosaic *Snrpb* deletion along the rostrocaudal axis at 36 and 48 hours post-tamoxifen injection.**
Representative fluorescence images of control and *Snrpb*$^{tmx+/–}$ mutant embryos carrying the *Rosa*26-mTmG reporter at 36 and 48 hours post-tamoxifen injection at E8.5, showing mosaic GFP expression along the rostrocaudal axis. FB, forebrain; MB, midbrain; HB, hindbrain; PM, presomitic mesoderm; TB, tailbud. Arrowheads indicate somites.

### Supplementary Figure 6.

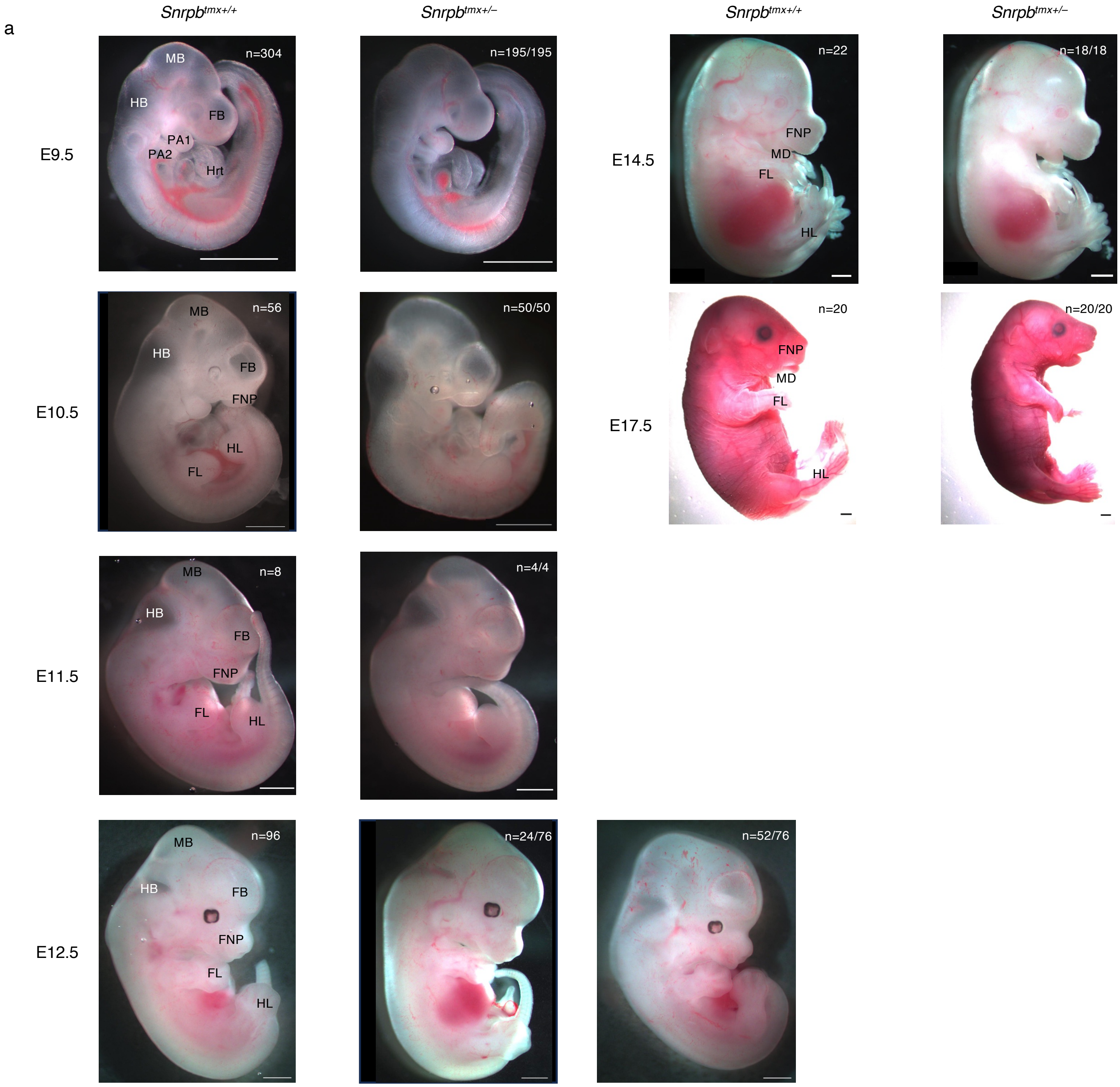

**Supplementary Figure 6.**

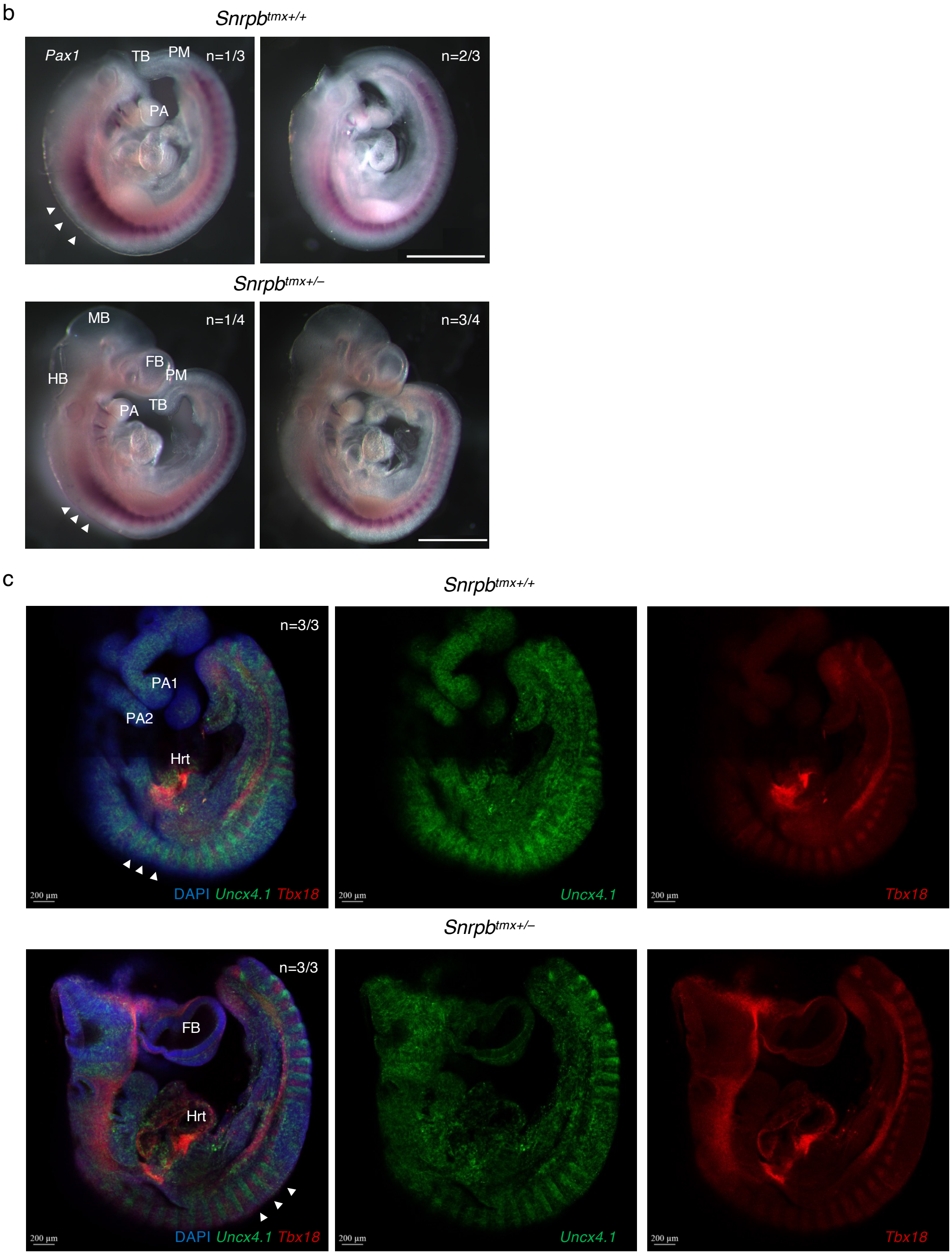


**Supplementary Figure 6. Morphological progression and rostrocaudal somite patterning in *Snrpb*$^{tmx+/-}$ mutant embryos.**

(a) Representative images of control and *Snrpb*$^{tmx+/-}$ mutant embryos collected between E9.5 and E17.5. Mutant embryos are morphologically indistinguishable from controls between E9.5 and E11.5. Morphological abnormalities first become apparent at E12.5, with approximately two-thirds of mutant embryos exhibiting a small, triangular forebrain, reduced frontonasal prominence, and shortened forelimb and hindlimb buds. At E14.5 and E17.5, mutant phenotypes are fully penetrant. Scale bars, 1mm. (b) Representative images of E9.5 control and *Snrpb*$^{tmx+/-}$ mutant embryos following whole mount *in situ* hybridization using probe targeting *Pax1*, sclerotome marker. Scale bars, 1mm. (c) Representative fluorescence images of E9.5 control and *Snrpb*$^{tmx+/-}$ mutant embryos following hybridization chain reaction (HCR) using probes targeting *Uncx4.1* and *Tbx18*, rostrocaudal polarity markers of the posterior and anterior somite compartments, respectively. Scale bars, 200$\mu$m. FB, forebrain; MB, midbrain; HB, hindbrain; PA1, pharyngeal arch 1; PA2, pharyngeal arch 2; Hrt, heart; FNP, frontonasal prominence; MD; mandible; FL, forelimb; HL, hindlimb; PM, presomitic mesoderm; TB, tailbud. Arrowheads indicate somites.

### Supplementary Figure 7.

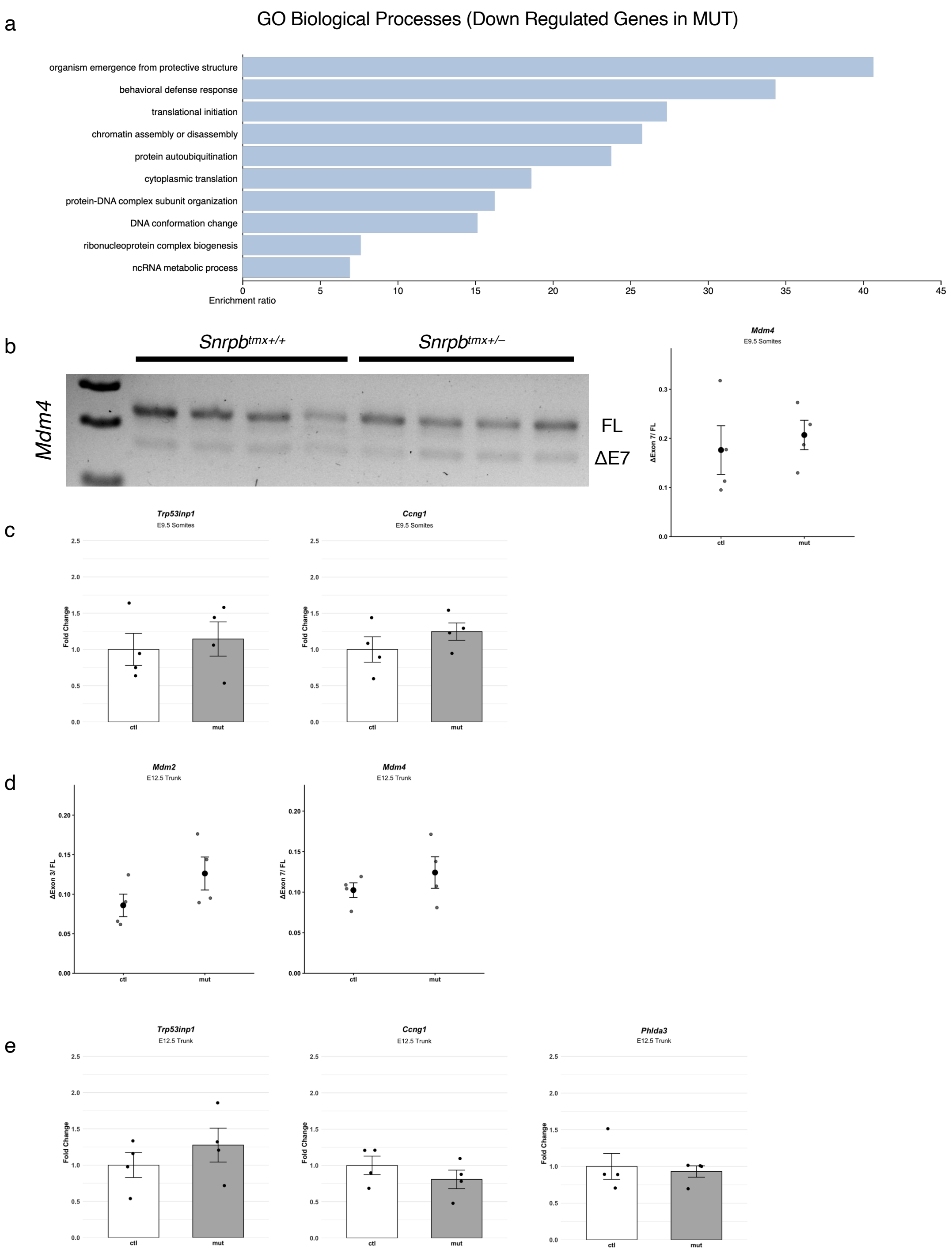


**Supplementary Figure 7. p53 pathway analysis in E9.5 *Snrpb*$^{tmx+/–}$ mutant somites and E12.5 trunk tissues.**

(a) Gene ontology biological process enrichment analysis of genes downregulated in *Snrpb*$^{tmx+/–}$ mutants relative to controls (FDR>0.05). (b) Representative RT-PCR gel showing alternative splicing of *Mdm4* exon 7 in E9.5 control and mutant somites, and dot plot (mean ± SEM) showing no significant change in *Mdm4* exon 7 skipping between genotypes (one-tailed Student's t-test, $p > 0.05$). (c) Bar graphs (mean ± SEM) showing *Trp53inp1* and *Phlda3* expression levels in E9.5 control and mutant samples, with no significant difference between genotypes. (d) Dot plots (mean ± SEM) showing *Mdm2* exon 3 skipping and *Mdm4* exon 7 skipping in E12.5 control and mutant trunk tissues. (e) Bar graphs (mean ± SEM) showing *Trp53inp1*, *Ccng1*, and *Phlda3* expression levels in E12.5 control and mutant trunk tissues, with no significant difference between genotypes. (f) Representative immunohistochemistry sections showing p53-positive cells (arrowheads) in posterior somites of E9.5 control and *Snrpb*$^{tmx+/–}$ mutant embryos, and dot plot (mean ± SEM) showing no significant change in p53-positive cell proportions in posterior somites between genotypes (one-tailed Student's t-test, $p > 0.05$).

### Supplementary Figure 8.

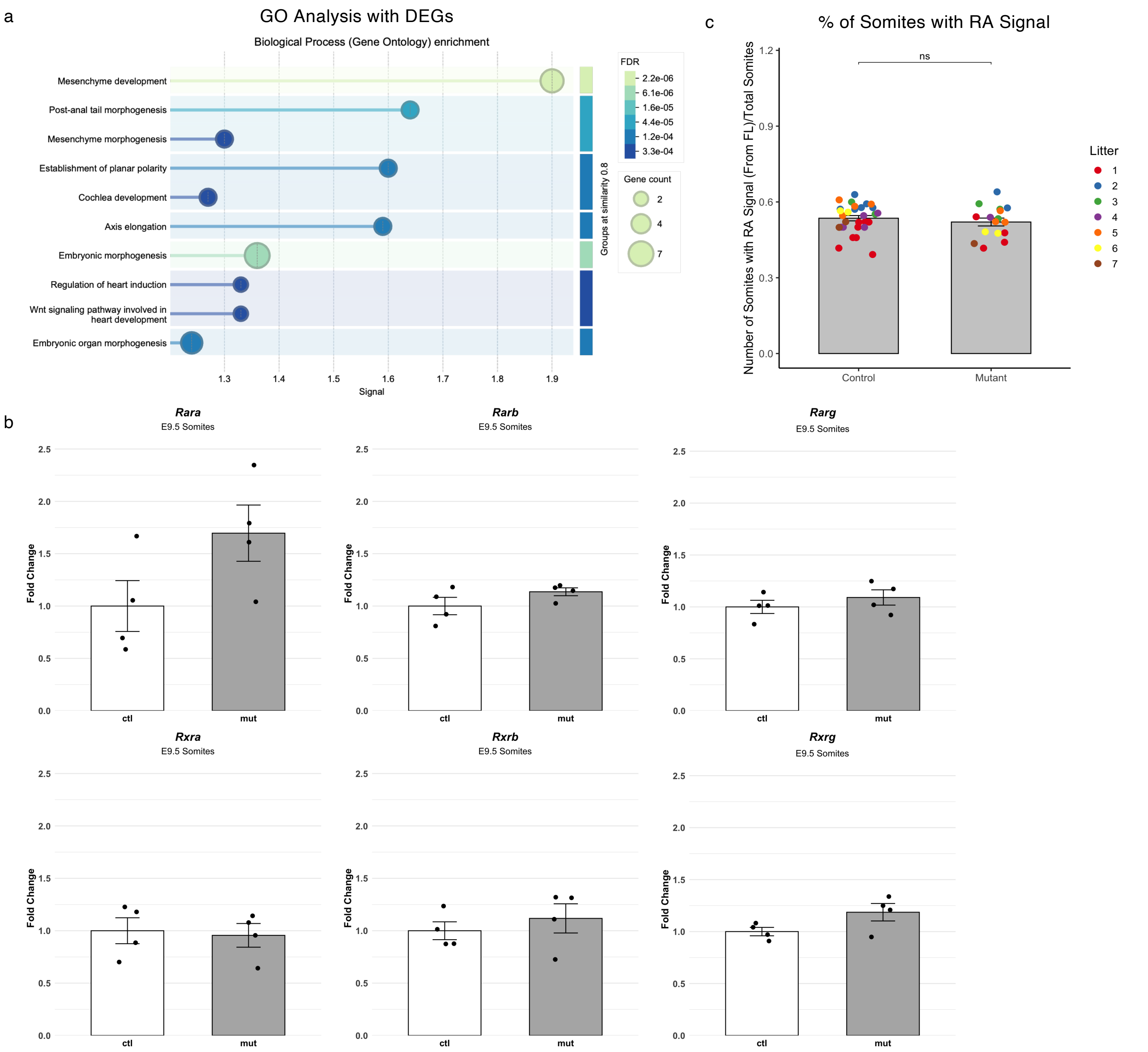


**Supplementary Figure 8. Pathway analysis of axial skeletal DEGs reveals mesenchyme development enrichment, with no change in RA receptor expression or spatial RA signaling domain in *Snrpb*$^{tmx+/–}$ mutant somites.**

(a) Gene ontology biological process enrichment analysis of differentially expressed genes (adjusted $p < 0.05$, fold change > 1) associated with rib and vertebral phenotypes in *Snrpb*$^{tmx+/–}$ mutants. (b) Bar graphs (mean ± SEM) showing RT-qPCR quantification of retinoic acid receptor (RAR) and retinoid X receptor (RXR) isoforms in E9.5 control and *Snrpb*$^{tmx+/–}$ mutant somites (one-tailed Student's t-test, $p > 0.05$). (c) Bar plots (mean ± SEM) showing the number of somites displaying LacZ signal from the forelimb region to the tailbud, and this value normalized to total somite number, in E9.5 control and *Snrpb*$^{tmx+/–}$ mutant embryos carrying the RARE–LacZ reporter (two-tailed Student's t-test, $p > 0.05$).

**Supplementary Figure 9.**

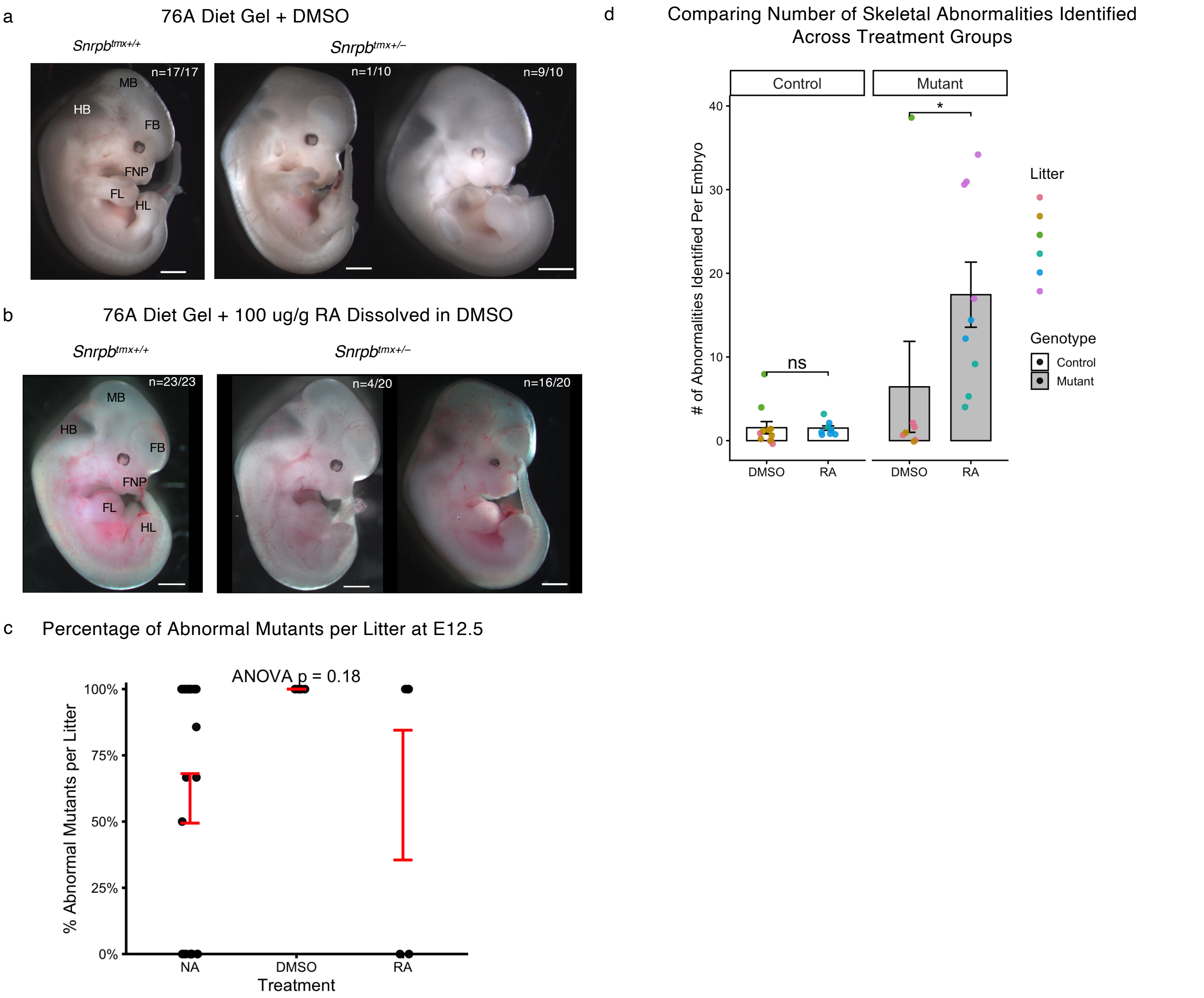


**Supplementary Figure 9. Effects of RA dietary supplementation on *Snrpb$^{tmx+/–}$* mutant embryos at E12.5 and at E18.5.**

(a) Representative images of E12.5 control and *Snrpb$^{tmx+/–}$* mutant embryos from dams fed 76A Diet Gel supplemented with DMSO from E7.5 to E9.5. All control embryos appeared morphologically normal. Among mutants, 9 out of 10 displayed the characteristic E12.5 phenotype with a triangular forebrain and reduced frontonasal prominence, and 1 out of 10 displayed a triangular forebrain with a split forelimb. (b) Representative images of E12.5 control and *Snrpb$^{tmx+/–}$* mutant embryos from dams fed 76A Diet Gel supplemented with 100 $\mu$g/g RA in DMSO from E7.5 to E9.5. All control embryos appeared morphologically normal. Among mutants, 4 out of 20 resembled control littermates, whereas 16 out of 20 displayed the characteristic E12.5 phenotype with a triangular forebrain and reduced frontonasal prominence. FB, forebrain; MB, midbrain; HB, hindbrain; FNP, frontonasal prominence; FL, forelimb; HL, hindlimb. Scale bars, 1mm. (c) Dot plot (mean ± SEM) comparing the proportion of morphologically abnormal mutant embryos per litter at E12.5 across treatment conditions (one-way ANOVA). (d) Bar plot (mean ± SEM) comparing the number of skeletal abnormalities identified per embryo at E18.5 across DMSO and RA treatment conditions. Individual data points represent embryos, color-coded by litter (two-way ANOVA with unadjusted pairwise comparisons of estimated marginal means between genotypes within each treatment; ns, not significant; *$p < 0.05$). NA, no treatment; DMSO, 76A Diet Gel + DMSO; RA, 76A Diet Gel + 100 $\mu$g/g RA in DMSO.

**Supplemental Figure 10.**

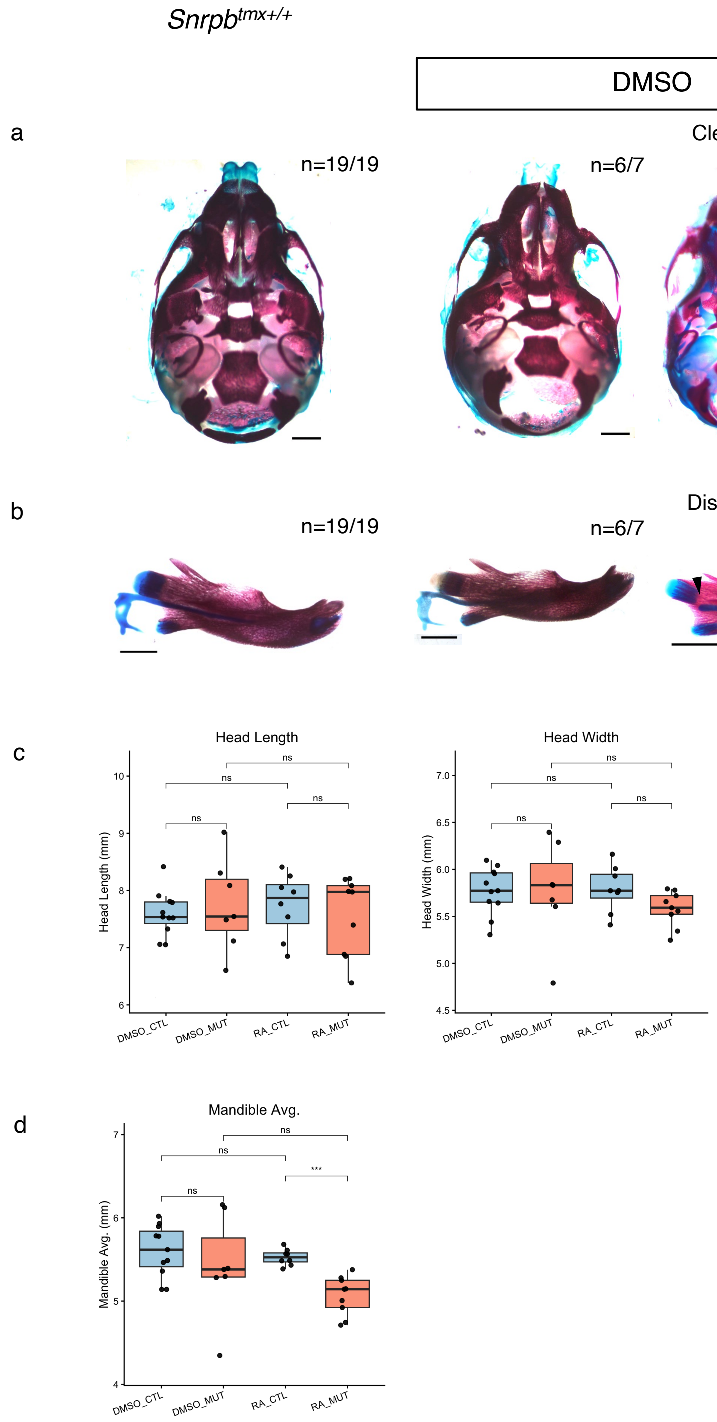


**Supplementary Figure 10. Craniofacial abnormalities in E18.5 *Snrpb$^{tmx+/–}$* mutants following maternal dietary RA supplementation.**

Representative skeletal preparations of E18.5 control and *Snrpb$^{tmx+/–}$* mutant embryos from dams fed 76A Diet Gel supplemented with DMSO or 100 $\mu$g/g RA in DMSO from E7.5 to E9.5, showing (a) cleft palate (arrowheads) and (b) discontinuous Meckel's cartilage (arrowheads). (c) Box plots comparing head length and head width and (d) averaged mandible length between control and *Snrpb$^{tmx+/–}$* mutant embryos within and across treatment groups (two-tailed Student's t-test; ***$p < 0.001$). Scale bars, 1mm.

**Supplementary Figure 11.**

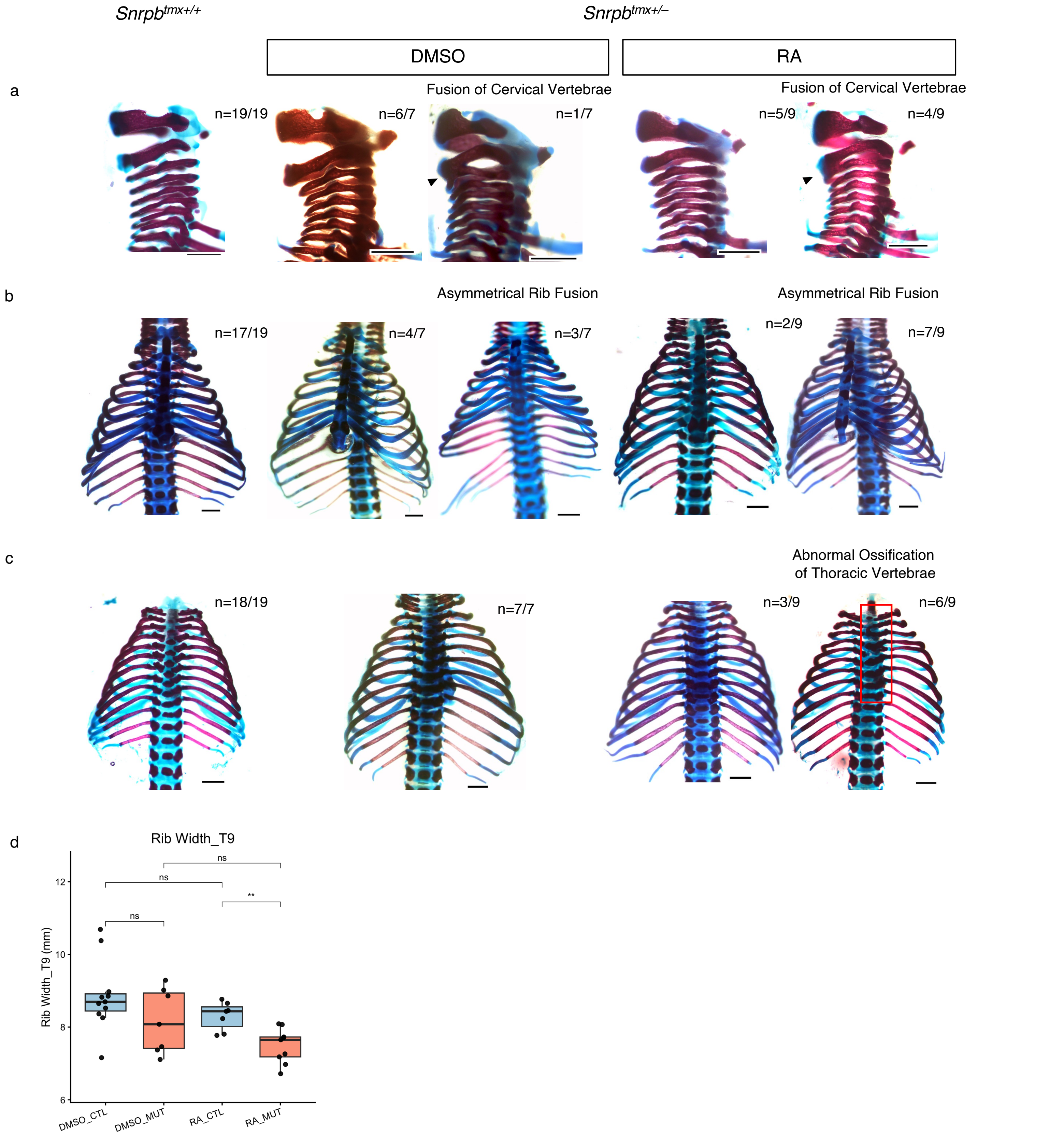


**Supplementary Figure 11. Rib and vertebral abnormalities in E18.5 *Snrpb*$^{tmx+/–}$ mutants following maternal dietary RA supplementation.**

Representative skeletal preparations of E18.5 control and *Snrpb*$^{tmx+/–}$ mutant embryos from dams fed 76A Diet Gel supplemented with DMSO or 100 $\mu$g/g RA in DMSO from E7.5 to E9.5, showing (a) cervical vertebral fusion (arrowheads), (b) asymmetric rib fusion, and (c) abnormal ossification of thoracic vertebrae (box). (d) Box plots comparing T9 rib width between control and *Snrpb*$^{tmx+/–}$ mutant embryos within and across treatment groups (two-tailed Student's t-test; **$p < 0.01$). Scale bars, 1mm.

**Supplementary Figure 12.**

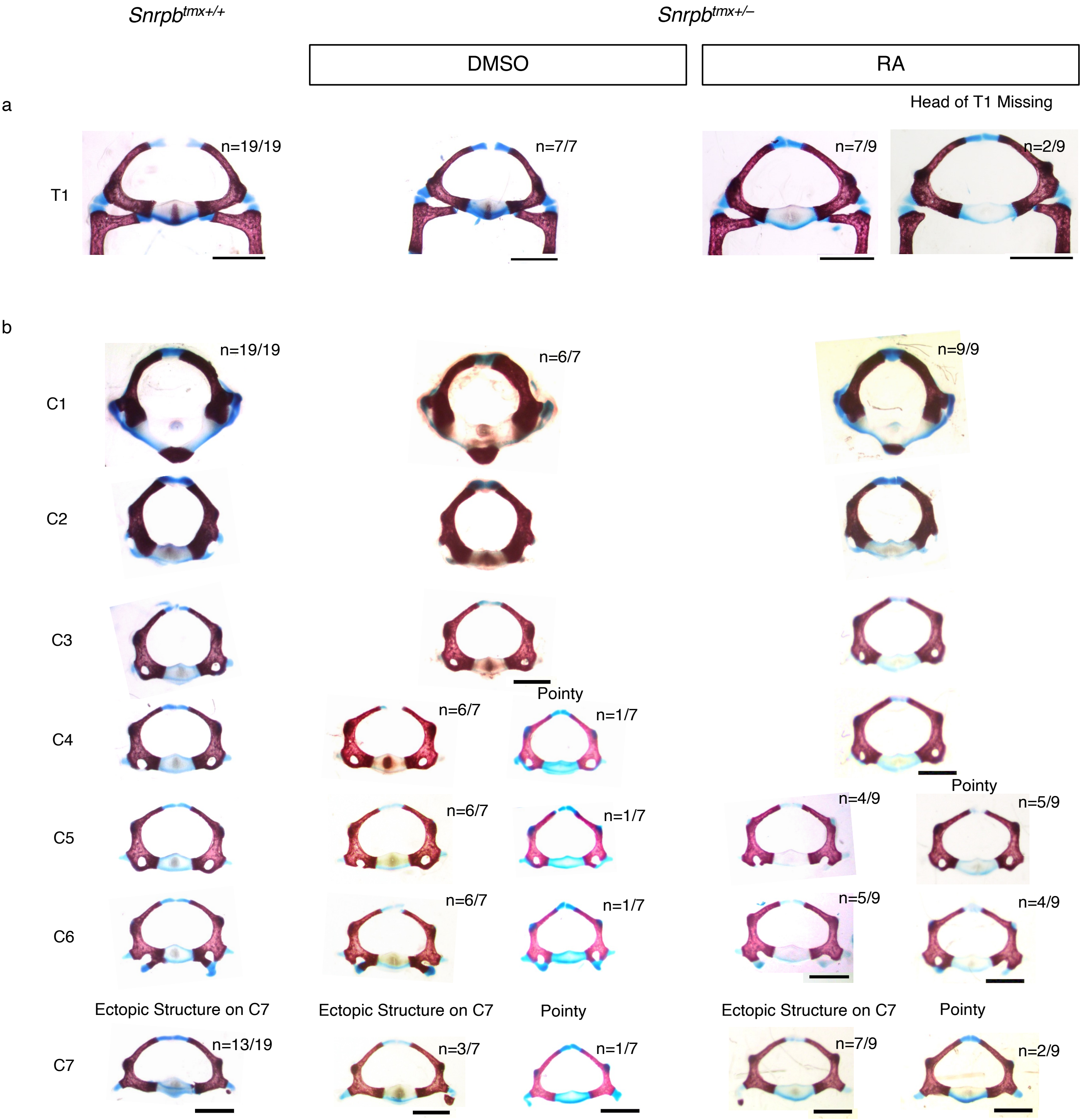


**Supplementary Figure 12. Thoracic and cervical vertebral abnormalities in E18.5 *Snrpb*$^{tmx+/–}$ mutants following maternal dietary RA supplementation.**

Representative skeletal preparations of E18.5 control and *Snrpb*$^{tmx+/–}$ mutant embryos from dams fed 76A Diet Gel supplemented with DMSO or 100 $\mu$g/g RA in DMSO from E7.5 to E9.5, showing (a) abnormal rib attachment to the first thoracic vertebra (T1) and (b) abnormal cervical vertebral morphology. Scale bars, 1mm.

### Supplementary Figure 13.

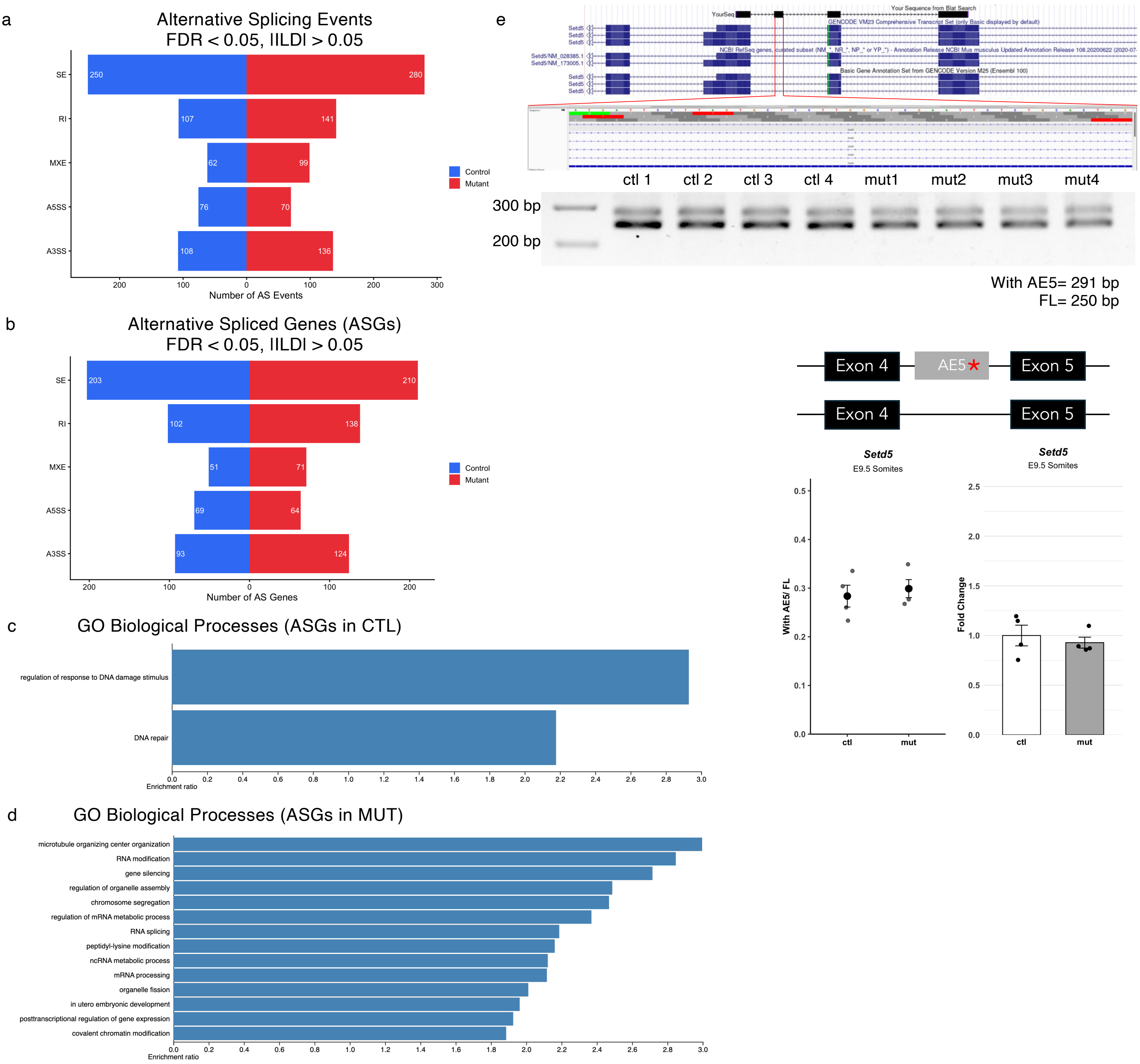


**Supplementary Figure 13. Alternative splicing landscape in E9.5 *Snrpb$^{tmx+/-}$* mutant somites and validation of *Setd5* alternative splicing.**

Diverging bar charts showing the number and distribution of (a) alternative splicing events (FDR < 0.05, IΙΔΙ > 0.05) and (b) alternatively spliced genes (FDR < 0.05, IΙΔΙ > 0.05) in E9.5 control and *Snrpb$^{tmx+/-}$* mutant somite samples. Gene ontology biological process enrichment analysis of genes alternatively spliced in (c) controls and (d) *Snrpb$^{tmx+/-}$* mutants. (e) Sanger sequencing confirming inclusion of alternative exon 5 (AE5) in *Setd5* introduces a premature stop codon (asterisk) in all three reading frames. Representative RT-PCR gel showing alternatively spliced *Setd5* isoforms in control and *Snrpb$^{tmx+/-}$* mutant somites, and dot plots and bar graphs (mean ± SEM) showing *Setd5* exon inclusion levels and transcript abundance by RT-PCR and RT-qPCR, respectively, with no significant difference between genotypes.
