## Supplementary Note for "Developmental pleiotropy revealed by mosaic heterozygous *Snrpb* deletion underlies CCMS-like axial skeletal defects"

**Determining the critical developmental window for tamoxifen-induced *Snrpb* deletion to model CCMS-like axial skeletal defects**

To investigate the role of ***Snrpb*** during embryonic development while bypassing early embryonic lethality associated with *Snrpb* insufficiency, we used a tamoxifen-inducible Cre–LoxP system to generate heterozygous deletion of ***Snrpb* at various embryonic stages (Table 1)**. Tamoxifen injection at **E8.0** resulted in early developmental abnormalities and reduced embryonic survival. By E11.5, mutant embryos (n = 6/8) displayed forebrain and midbrain hypoplasia and a laterally expanded cephalic flexure. They also lacked the distal paddle-shaped structures of the forelimbs (Supplementary Figure 1A; Supplementary Data 1). At E12.5, mutants were recovered at significantly lower than expected Mendelian ratios (Supplementary Figure 1A; Supplementary Data 1). Together, these findings indicate that proper *Snrpb* levels are required for early embryonic survival, thereby preventing analysis of later skeletal phenotypes and prompting us to examine later induction timepoints.

To allow embryos to survive to later stages for phenotypic analysis, we next induced Snrpb deletion between E8.5 and E9.5 and assessed embryos at E14.5. In contrast to induction at earlier time points, deletion within this window permitted survival to E14.5 at expected Mendelian ratios (Table 1; Supplementary Data 1) but did not result in CCMS-like axial skeletal abnormalities. Mutant embryos appeared comparable to controls at time of collection, and cartilage preparations revealed normal craniofacial structures as well as rib and vertebral patterning (Supplementary Figure 1B-C). Although a reduction in head length was observed, measurements of Meckel’s cartilage and rib length were not significantly altered (Supplementary Figure 1D-E). These results indicate that Snrpb deletion between E8.5 and E9.5 is insufficient to disrupt rib and vertebral development at E14.5, thereby narrowing the critical developmental window for disease modeling.
